# Coordination Between Embryo Growth and Trophoblast Migration Upon Implantation Delineates Mouse Embryogenesis

**DOI:** 10.1101/2022.06.13.495767

**Authors:** V. Bondarenko, M. Nikolaev, D. Kromm, R. Belousov, A. Wolny, S. Rezakhani, J. Hugger, V. Uhlmann, L. Hufnagel, A. Kreshuk, J. Ellenberg, A. Erzberger, M. Lutolf, T. Hiiragi

**Author notes:** Hubrecht Institute, Uppsalalaan 8, 3584 CT Utrecht, Netherlands (V.B., T.H.); Institute for Translational Bioengineering (ITB), Roche Pharma Research and Early Development, CH-4070 Basel, Switzerland (M.N., M.L.); Novartis Institutes for BioMedical Research, Novartis Pharma AG, 4056 Basel, Switzerland (S.R); Delft Center for Systems and Control, Delft University of Technology, Delft, Netherlands (D.K.); Veraxa Biotech, Heidelberg, Germany (L.H.). Correspondence (A.E.), (M.L.), (T.H.).

## Abstract

Implantation marks a key transition in mammalian development. The role of embryo-uterus interaction in periimplantation development is however poorly understood due to inaccessibility *in utero.* Here, we develop an engineered uterus-like microenvironment to recapitulate mouse development *ex vivo* up to E5.25 and discover an essential role of integrin-mediated trophoblast adhesion to the uterine matrix. Light-sheet microscopy shows that trophoblast cells undergo Rac1-dependent collective migration upon implantation, displacing Reichert’s membrane and generating space for egg cylinder growth. The key role of coordination between trophoblast migration and embryo growth is verified by experimentally manipulating the migration velocity and geometry of the engineered uterus. Modeling the implanting embryo as a wetting droplet links the tissue shape dynamics to underlying changes in trophoblast adhesion and suggests that the corresponding tension release facilitates egg cylinder formation. Together, this study provides mechanisms by which dynamic embryo-uterus interactions play an essential role in peri-implantation development.

## INTRODUCTION

Organismal development is coordinated in space and time across multiple scales. Spatial coordination of mechanics and cell proliferation underlies tissue growth and morphogenesis (Wang and Riechmann, 2007; Lecuit and le Goff, 2007), and the spatial context can determine stem-cell fate (Rompolas et al., 2013). Oscillatory gene expression is linked to tissue-level morphogenesis and patterning leading to vertebrate segmentation (Palmeirim I et al., 1997). Spatial coordination can go beyond embryos – extra-embryonic tissues regulate the growth and patterning of embryonic tissues via mechano-chemical signaling (Brennan J. et al., 2001; Chen S-R. and Kimelman D., 2000). Furthermore, the maternally provided vitelline envelope physically interacts with embryonic tissues to coordinate their movement (Bailles et al., 2019; Münster et al., 2019). However, studying the interplay between the triad of embryonic – extraembryonic – maternal tissues in viviparous mammals remains a challenge and its potential role is poorly understood.

By establishing the embryo-maternal interaction, implantation represents a critical developmental stage in mammalian species. Mammalian development begins with generating extra-embryonic lineages, trophectoderm (TE) and primitive endoderm (PrE), in addition to the embryonic epiblast (EPI) in the blastocyst. In mice, at embryonic day (E) 4.5, the TE differentiates into EPI-attaching polar TE (pTE) and EPI-distant mural TE (mTE), which adheres to the uterine wall and initiates implantation. The former generates the extraembryonic ectoderm (ExE) tissue and the latter differentiates into giant trophoblast (GT), while EPI and ExE proliferate and elongate to form an “egg cylinder”. The extraembryonic lineages neighboring the EPI, ExE and visceral endoderm (VE) derived from PrE, play a key role in embryonic growth, patterning and body-axis formation via signaling (Brennan J. et al., 2001; Ichikawa et al., 2022; Rodriguez et al., 2005). In addition to these interactions between embryonic and extra-embryonic tissues post-implantation, the maternal uterine tissues establish a unique context for mammalian development. In particular, while the placenta provides nutritional support for embryonic growth post-implanation, the potential role of embryo-uterus interactions in periimplantation embryogenesis remains largely unexplored (Hiramatsu et al., 2013; Mesnard et al., 2004), as this process is inaccessible *in utero* and none of the *ex vivo* culture available to date allows successful reproduction of tissues mediating the embryo-uterus interaction.

*Ex vivo* culture of peri-implantation mouse embryos has been developed both in 2D (Hsu, 1972, 1973; Morris et al., 2012; Bedzhov et al., 2014) and 3D (Govindasamy et al., 2021; Ichikawa et al., 2022). However, the 2D culture disrupts morphogenesis as embryos adhere to the 2D surface, and the 3D culture so far has required removal of the mTE to release tension in the pTE enabling invagination and formation of the ExE. While our recent data suggested the role of embryo-uterus interaction in tension release *in utero* (Ichikawa et al., 2022), the exact mechanism awaited development of an *ex vivo* system that recapitulates the uterine environment and embryo-uterus interaction upon implantation.

Bioengineering approaches are generally used to emulate biochemical and mechanical properties of *in vivo* tissues and identify microenvironmental characteristics necessary and/or sufficient for tissue morphogenesis and patterning. For example, biomaterials engineering and microfabrication provide powerful tools for such *ex vivo* modeling of the native 3D environment (Lutolf and Hubbell, 2005; Vianello and Lutolf, 2019). Chemically-defined matrices, such as those based on poly(ethylene glycol) (PEG), provide tunability, robustness, and reproducibility for state-of-the-art mechanobiological studies (Seliktar, 2012; Caliari and Burdick, 2016; Gjorevski et al., 2016; Qazi et al., 2022). These techniques can be combined with live microscopy to gain mechanistic insights into tissue morphogenesis and patterning (Gjorevski et al., 2022).

Building upon these recent developments, in this study we develop an engineered uterus that reconstitutes key properties of the mouse uterine environment at implantation, in order to dissect the potential role and mechanisms of embryo-uterus interaction and tissue coordination during mouse peri-implantation development.

## RESULTS

### An engineered uterus recapitulates embryo-uterus interaction *ex vivo* and supports peri-implantation development of the whole embryo

Based on our recent findings (Ichikawa et al., 2022), we reasoned that *ex vivo* development of the whole embryo with ExE and Reichert’s membrane would require recapitulating embryo-uterus interaction *in utero.* Such an *ex vivo* system would be a prerequisite to studying the role of embryo-uterine interaction. To this end, we aimed to engineer a fully controllable uterine environment that mimics key biochemical and mechanical properties of the native uterine tissue. With this goal, we first analyzed the deposition of the extra-cellular matrix (ECM) at the embryo implantation sites of the pregnant uteri at embryonic day 4.75 (E4.75) and E5.25. Major ECM components, including fibronectin, collagen IV, and laminin, surround the elongated embryos (Figure 1A, B, Figure S1A) (Farrar and Carson, 1992).

**Figure 1.**
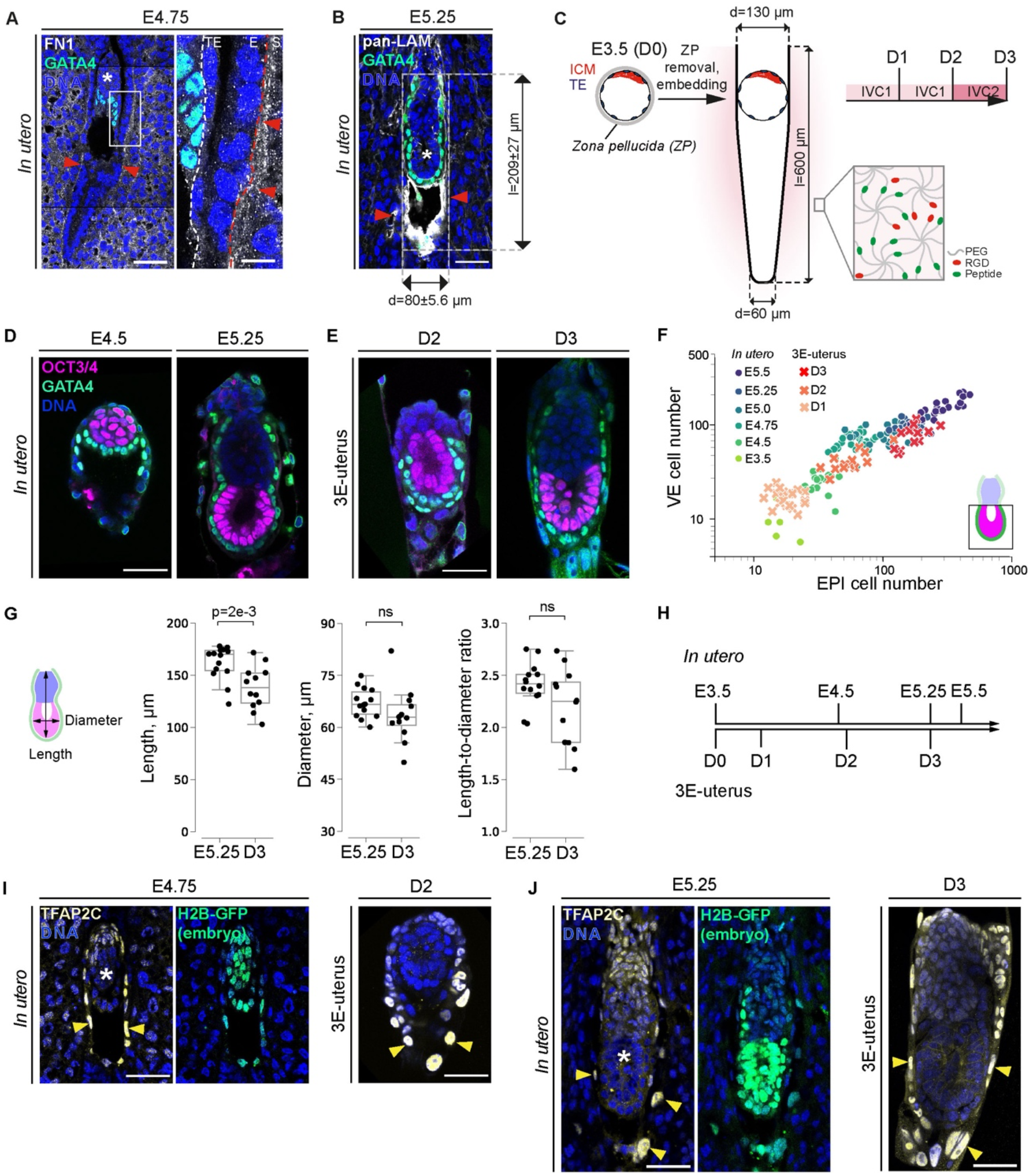
An *ex vivo* culture using 3D patterned hydrogel recapitulates the uterine mechano-chemical environment for peri-implantation mouse development. **(A)** Left, representative immunofluorescence image of the E4.75 uterus cross-section, simultaneously stained for Fibronectin (FN1, white), GATA4 (green), and nuclei (DNA, blue). Right, 4x zoom into the interface between trophectoderm (TE), uterine epithelium (E), and stroma (S) (outlined). Red arrowheads mark uterine ECM. **(B)** Representative immunofluorescence image of the E5.25 uterus cross-section, simultaneously stained for pan-Laminin (panLAM, white), GATA4 (green), and nuclei (DNA, blue). Measurements of E5.25 *in utero* embryo dimension across n = 12. **(C)** Schematic illustration of peri-implantation embryo culture. Embryos are recovered at E3.5, treated by Tyrode’s solution to remove *Zona pellucida* (ZP), and embedded into the crypts on the day of recovery (D0). ICM, red; TE, blue; IVC1 and ICV2 stand for “In Vitro Culture” medium 1 and 2, respectively. Inset, schematic illustration of the hydrogel composition. 8-arm Poly(ethylene glycol) (PEG, gray) molecules, crosslinked via metalloprotease-cleavable peptides (Peptide, green), and functionalized with RGDSPG peptide (Arg-Gly-Asp-Ser-Pro-Gly, ‘RGD’, red). **(D)** Representative immunofluorescence images of the embryos developed *in utero* until E4.5 and E5.25, simultaneously stained for OCT3/4 (magenta), GATA4 (green), and nuclei (DAPI, blue). **(E)** Representative immunofluorescence images of the 3E-uterus embryos from day 2 and 3, simultaneously stained for OCT3/4 (magenta), GATA4 (green), and nuclei (DAPI, blue). **(F)** Scatterplot showing numbers of epiblast (EPI) cells (x-axis) vs numbers of visceral endoderm (VE) cells (y-axis) that cover epiblast in the embryos developed *in utero* until E5.5 (E3.5 – E5.5) and the embryos developed by 3E-uterus until day 3 (D1 – 3). n = 5 (E3.5), n = 21 (E4.5), n = 28 (E4.75), n = 20 (E5.0), n = 20 (E5.25), n = 21 (E5.5), data from (Ichikawa et al., 2022); n = 20, two replicates pooled (D1), n = 13 of 28, three replicates pooled (D2), n = 12 of 26, three replicates pooled (D3). X/Y scale, log 10. Bottom right, the scheme of the quantified embryo region, EPI (magenta), VE (green). **(G)** From left to right, boxplots showing egg cylinder’s length, diameter, and the length-to-diameter ratio between embryos developed *in utero* until E5.25 and 3E-uterus embryos from day 3 (D3). n = 14 and 12, respectively. Data points, shown as black dots, correspond to individual embryos, midline marks the median, boxes indicate interquartile range. P-values were calculated using Student’s t-test and the Mann-Whitney U test. **(H)** Cell number-based correspondence between *in utero* and 3E-uterus embryo development. **(I)** Left, representative immunofluorescence image of the E4.75 uterus cross-section from the pregnant F1 female mated with the H2B-GFP male. Simultaneous immunostaining for TFAP2C (yellow), GFP (green), and nuclei (DNA, blue). Right, immunostaining of the 3E-uterus embryo from day 2 (D2). Yellow arrowheads mark differentiated trophoblast cells. **(J)** Left, representative immunofluorescence image of the E5.25 uterus cross-section from the F1 female mated with the H2B-GFP male. Simultaneous immunostaining for TFAP2C (yellow), GFP (green), and nuclei (DNA, blue). Right, immunostaining of the 3E-uterus embryo from day 3 (D3). Scale bars, 50 μm, 12.5 μm (4x zoom). White asterisks mark epiblast of the implanted embryos. See also Materials and Methods, Figure S1.

Accordingly, we first replaced hybrid Matrigel/collagen matrices in our recent 3D-culture, 3D-geec (Ichikawa et al., 2022), with synthetic poly(ethylene glycol) (PEG)-based hydrogels. To mimic the uterine biochemical characteristics, LDTM PEG gels (Low-Defect Thiol-Michael addition hydrogels, thereafter referred to as hydrogel) (Rezakhani et al., 2020) were functionalized with the fibronectin-derived, integrin-binding adhesion ligand RGDSPG (Arg-Gly-Asp-Ser-Pro-Gly, ‘RGD’), and cross-linked via matrix metalloprotease (MMP)-cleavable peptide substrates (Figure 1C, inset; Figure S1B). However, E3.5 blastocysts embedded into 3D hydrogel drops failed to develop further, displaying growth retardation and impaired morphogenesis as judged by significantly fewer epiblast cells than in the periimplantation embryos developed *in utero* (Figure S1C, D). Notably, in these synthetic isotropic 3D culture microenvironments, the growth of the epiblast tissue appears to be blocked by the presence of Reichert’s membrane (Figure S1C; Ichikawa et al., 2022). These observations suggest that an isotropic gel environment fails to recapitulate key embryo-uterine interactions and point to the additional need of emulating geometrical context and mechanical properties of the native uterine tissue.

Thus, using microfabrication, we applied a topographical 3D modification of the hydrogel to generate with high precision (Nikolaev et al., 2020; Gjorevski et al., 2022) the elongated crypt shape that mouse uteri acquire around the implanting embryo (Burckhard, 1901; Cha et al., 2014; Figure 1C, Video S1). The optimal crypt dimension was determined by the efficiency of the *ex vivo* culture, as judged by the embryo morphology (Figure S1E, F). A diameter gradient was introduced to accommodate variability in blastocyst size (Figure 1C). Among the 10-fold range of the shear moduli generated by 1.5-7% PEG precursor content, 1.5-2 % PEG generated the shear modulus at 100-300 Pa, which is in the stiffness range of the E5.5 mouse decidua (Govindasamy et al., 2021) and resulted in the highest developmental efficiency (Figure S1G, H).

Under these conditions, E3.5 blastocysts developed *ex vivo* for 3 days (D3) and reached epiblast (EPI) and visceral endoderm (VE) cell numbers comparable to E5.25 embryos developed *in utero* (Figure 1D-F, Video S1). Embryo diameter and length-to-diameter ratio were comparable, although length itself seems shorter after *ex vivo* culture (Figure 1G). Based on the cell numbers, the temporal developmental progression is estimated to be slowed down for the first two days of *ex vivo* culture, with a total delay of 30 hours (Figure 1H, Figure S1I). Overall, our engineered uterus system reproduces E5.25 egg cylinder formation (Figure 1D-H) with a 46% efficiency (n = 12 of 26 embryos) after 3 days of culture.

Laminin-rich Reichert’s membrane, connected to the basal membrane of the egg cylinder, successfully formed in 77% of the embryos (n = 20 of 26 embryos) (Figure S1I, J). In these embryos, the inner side of the Reichert’s membrane contained GATA4-positive cells, corresponding to the parietal endoderm (PE), whereas Tfap2C- and Krt8-positive trophoblast (TB) formed on the outer side with enlarged nuclei (Figure 1J and Figure S1J).

Altogether, these results indicate that our new method of *ex vivo* culture with an engineered uterus, named 3E-uterus (*<u>e</u>x vivo* <u>e</u>ngineered uterine <u>e</u>nvironment with 3D geometrically patterned hydrogels), supports differentiation of embryonic as well as extraembryonic tissues, closely recapitulating *in utero* peri-implantation mouse development.

### Adhesion to the uterine matrix is essential for peri-implantation mouse development

As the geometrical and mechanical context for the engineered uterus is established, we examined the dependence of *ex vivo* embryo development on its key biochemical characteristics. Strikingly, the developmental efficiency significantly dropped in the absence of RGD (Figure 2A, B), suggesting that integrin-mediated adhesion of the embryo to the uterine wall is absolutely required for peri-implantation mouse development. In agreement with this, the integrin beta 1 subunit and its active form are enriched at the basal as well as the apical side of the mTE/TB cells that mediate adhesion of the embryo to the uterine wall (Figure 2C; A. E. Sutherland, 1993; Govindasamy et al., 2021), in contrast to its basal localization in E3.5 blastocysts (Figure S2A; Kim et al., 2022).

**Figure 2.**
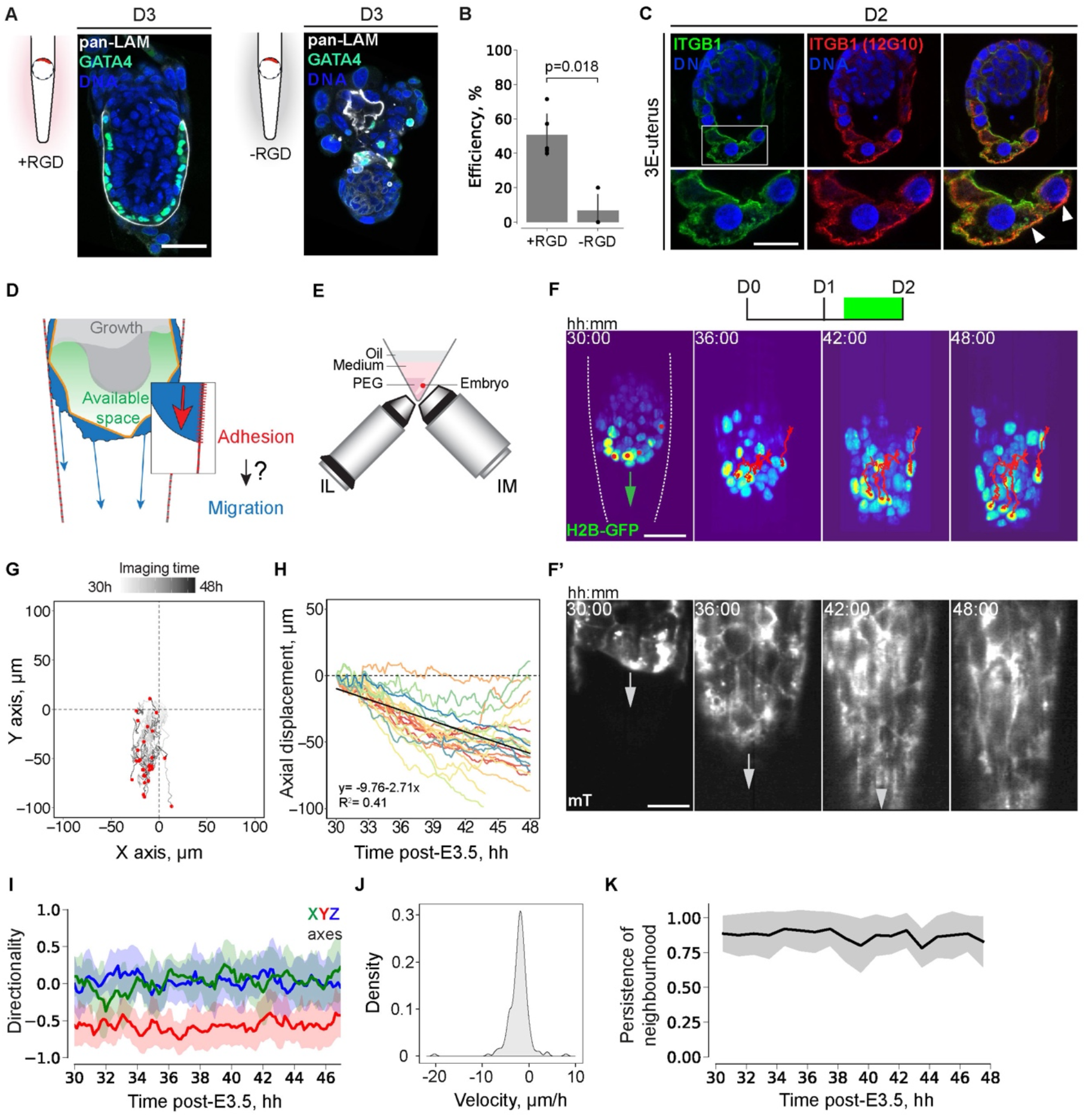
Trophoblast cells adhesion to the uterine matrix triggers their collective migration and is essential for periimplantation mouse development. **(A)** Left, representative immunofluorescence image of the embryo cultured in the crypts made of PEG with RGD for 3 days and stained for pan-Laminin (pan-LAM, white), GATA4 (green), and nuclei (DNA, blue). Right, representative immunofluorescence image of the embryo cultured in the crypts made of PEG without RGD for 3 days. **(B)** Bar-plots showing 3E-uterus developmental efficiency with and without RGD. n = 5 and 3, respectively. Dots correspond to efficiency values in experimental replicates, whiskers indicate standard deviations. P-values were calculated using Student’s t-test. **(C)** Representative immunofluorescence image of the embryos cultured in the crypts made of PEG with RGD for 2 days, simultaneously stained for total integrin beta 1 (ITGB1, green), active ITGB1 (12G10, red), and nuclei (DNA, blue). From left to right, total ITGB1, active ITGB1, composite image channels. Bottom, 2x zoom. White arrowheads mark the apical surface of the trophoblast cells. **(D)** Schematic illustration of the hypothesis that adhesion-induced collective migration of TB cells generates space for embryo growth. **(E)** Schematic for the inverted light-sheet microscopy with illumination objective (IL) and imaging objective (IM). **(F)** Representative 3D projections of time-lapse images of the H2B-GFP in the embryo developing in the crypts made of PEG with RGD. Trajectories of individual mural TE cells are marked with red lines. The crypt surface is outlined. t =00:00, hours: minutes after recovery at E3.5. See also Video S2. **(F’)** Representative time-lapse images of the same embryo, showing the plasma-membrane mTomato signal (grey) in the image plane corresponding to the crypt surface. **(G)** Trajectories of mural TE cells in an XY plane, normalized to the starting coordinates. End coordinates are marked with red dots. n = 29. **(H)** Displacement of mural TE cells along the Y-axis in relation to imaging time. n = 29. The linear regression fit is shown in black, y = −9.76-2.71x, R^2^ = 0.41. **(I)** Directionality of the mTE/TB migration along the X, Y, and Z axes (green, red, and blue, respectively) between subsequent time points (every hour of live imaging). n = 29. **(J)** Distribution density of the average TB velocities (μm/h). n = 255, pooled from 6 embryos. **(K)** Persistence of the nearest mTE/TB four-cell neighbourhood between consecutive time points (every hour of live imaging), n = 29. Scale bars, 50 μm, 25 μm (2x zoom), 12.5 μm (4x zoom). See also Figure S2, Materials and Methods.

Taken together, this engineering approach identified mechano-chemical properties of the uterine tissue that underlie the embryo-uterine interaction upon implantation. In particular, our findings suggest that adhesion of the embryo to the uterine matrix is necessary for peri-implantation mouse development.

### Trophoblast cells adhesion to the uterine matrix triggers their collective migration

We then investigated the mechanisms by which adhesion of the embryo to the uterine matrix plays an essential role in peri-implantation mouse development. Given the phenotype of the failure resulting from the 3D isotropic culture (Figure S1C; Ichikawa et al., 2022) and redistribution of integrin beta 1 to the apical and adhesion side of mTE/TB cells (Figure 2C), we hypothesized that mTE/TB cells may migrate upon adhesion to the uterine matrix and displace Reichert’s membrane to accommodate epiblast tissue growth (Figure 2D). To monitor mTE/TB cell dynamics upon adhesion to the uterine matrix, we integrated our hydrogel culture with inverted light-sheet microscopy (InVi-SPIM; Figure 2E; Strnad et al., 2016; Ichikawa et al., 2022). Tracking mTE/TB cell nuclei labeled with H2B-GFP (Hadjantonakis and Papaioannou, 2004) and the plasma membrane labeled with mTomato indeed showed cell migration along the crypt surface (Figure 2F, Video S2). Individual mTE/TB cells preferentially migrated downward along the crypt axis (Figure 2G-I and Figure S2B-C) with an average velocity of 2.51 μm/h (Figure 2J; Figure S2D) and maintaining the nearest neighbors (Figure 2K, Figure S2E). Nuclear division was rarely observed (12% of mTE/TB cell nuclei divided during 24 hours of live-imaging; n=158), suggesting a limited contribution of mTE/TB cell division to their displacement. Together, these results indicate that upon integrin-mediated adhesion, mTE/TB cells undergo collective migration.

### Trophoblast cells lose polarity and acquire mesenchymal motility upon adhesion

To understand the mechanisms of mTE cell reaction upon implantation, we further characterized mTE/TB cellular dynamics at the sub-cellular level. First, we examined how the apical side of mTE cells, which initially lacks integrin beta 1 (see Figure S2A), could adhere and mediate migration in the uterine ECM. Immunofluorescence staining of 3E-uterus embryos at D2 showed localization of the apical marker, pERM, at the basolateral surface and of the basal marker, integrin beta 1, at the apical surface (Figure 3A-D; see also Figure 2C). Localization of the tight-junction marker, ZO-1, also becomes disorganized during 3E-uterus culture (Figure 3A-D, Figure S3A-C, Video S3). These data show that mTE cells lose cell polarity upon adhesion to the uterine matrix and acquire a mesenchymal property.

**Figure 3.**
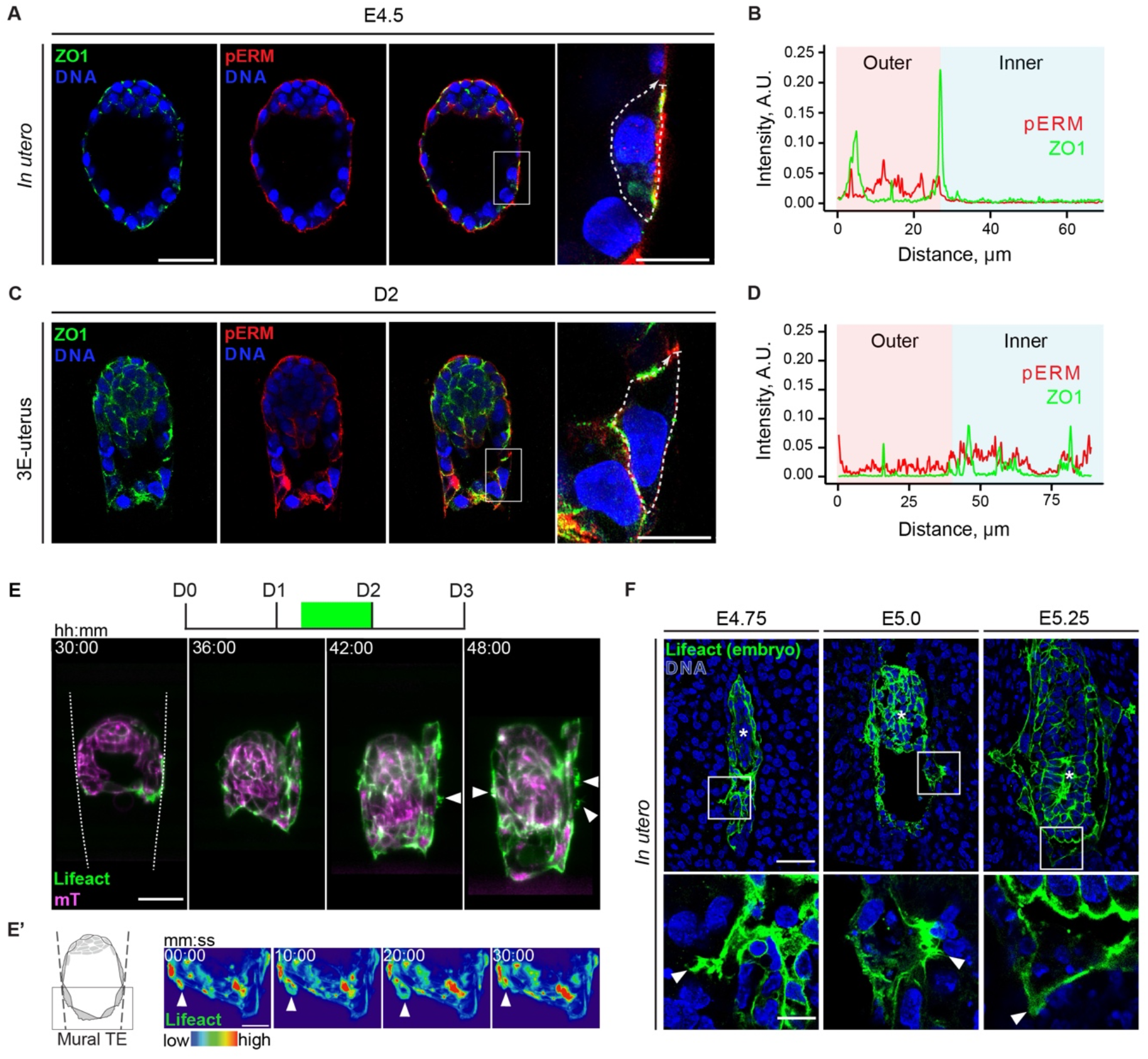
Trophoblast cells lose polarity and acquire mesenchymal motility upon adhesion. **(A)** Representative immunofluorescence image of the E4.5 embryo developed *in utero,* simultaneously stained for ZO-1 (green), phospho-Ezrin/Radixin/Moesin (pERM, red), and nuclei (DNA, blue). From left to right, ZO1, pERM, composite image channels. Right, 4x zoom into a mTE cell. **(B)** Intensity profile for ZO1 and pERM signals along the cell surface outlined in A (right), including apical and basolateral regions. **(C)** Representative immunofluorescence image of the 3E-uterus embryo from day 2, simultaneously stained for ZO-1 (green), phospho-Ezrin/Radixin/Moesin (pERM, red), and nuclei (DNA, blue). From left to right, ZO1, pERM, composite image channels. n = 5. Right, 4x zoom into a TB cell. **(D)** Intensity profile of ZO1 and pERM signals along the cell surface outlined in C (right), including apical and basolateral regions. **(E)** Representative time-lapse images of the developing Lifeact-GFP;mTmG embryo; Lifeact-GFP (green), mTomato (magenta). The crypt surface is outlined. White arrowheads point at the trophoblast membrane protrusions. t=00:00, hours: minutes after recovery at E3.5. See also Video S4. **(E’)** Representative 3D projection time-lapse images of mural TE of Lifeact-GFP in the developing embryo; Lifeact-GFP (spectral). White arrowheads point at lamellipodia. t=00:00, minutes: seconds. See also Video S5. **(F)** Left to right, representative immunofluorescence images of the E4.75, E5.0, and E5.25 uterus cross-sections from F1 females mated with Lifeact-GFP males; simultaneous staining for GFP (green) and nuclei (DNA, blue). Bottom, 4x zoom. Scale bars, 50 μm, 20 μm (E’), 15 μm (A, C, right), 12.5 μm (F, 4x zoom). See also Figure S3.

In line with this, light-sheet microscopy of actomyosin dynamics in Lifeact-GFP;mTmG and Myh9-GFP;mTmG (Zhang et al., 2012) embryos (Videos S4-6) showed enrichment at the adhesion side of mTE, followed by lamellipodia and filopodia formation at the TB migration front along the crypt axis and laterally (Figure 3E, Figure S3D, E). These data demonstrate that adhesion to the uterine matrix induces mTE/TB cells to lose polarity and acquire mesenchymal motility.

To investigate whether mTE/TB cells undergo migration upon implantation *in utero,* we systematically examined mTE/TB cells in their native uterine tissue context throughout the implantation stages. To distinguish embryo-derived cells in the uterine tissues, we crossed Lifeact-GFP (Riedl et al., 2010) males with WT females, so that only embryo-derived cells have GFP expression within the tissue sections. mTE/TB cells of the embryo formed actin-rich, filopodia/lamellipodial cell membrane protrusions into the uterine tissue at E4.75 (Figure 3F left). TB cell protrusions were observed along the mesometrial/anti-mesometrial (M/AM) axis, as well as laterally at E5.0 (Figure 3F, middle). At E5.25, TB cells protrusion extended 26.9±8.6 μm along the M/AM axis and 17.9±5.9 μm laterally (Figure 3F, right). These prominent filopodia/lamellipodia are in agreement with the migratory activity of mTE/TB cells *in utero*.

Altogether, these data strongly suggest that upon implantation, adhesion to the uterine matrix triggers mTE/TB cells to lose polarity and acquire mesenchymal motility, leading to their collective migration along the surface and into the uterine tissues (Bevilacqua and Abrahamsohn, 1988; Sutherland, 2003).

### A droplet-wetting process can explain embryo-uterus tissue-tissue interactions upon implantation

Next, to characterize the tissue-scale change upon implantation at a cellular resolution, we combined 3E-uterus with the multi-view light-sheet microscopy (MuVi-SPIM) (Krzic et al., 2012; McDole et al., 2018) by implementing a controlled environment to the sample chamber (Figure 4A). Live-imaging showed distinct changes in the position and contact angle of the embryo in relation to the 3E-uterus surface (Figure 4B). Our findings on the enrichment of integrins in TB cells (see Figures 2C, S2A) led us to a hypothesis that an active adaptation of embryo-uterus adhesion may explain the observed evolution of the embryo shape as described by the physics of a droplet wetting process (de Gennes 1985, Douezan et al. 2011).

**Figure 4.**
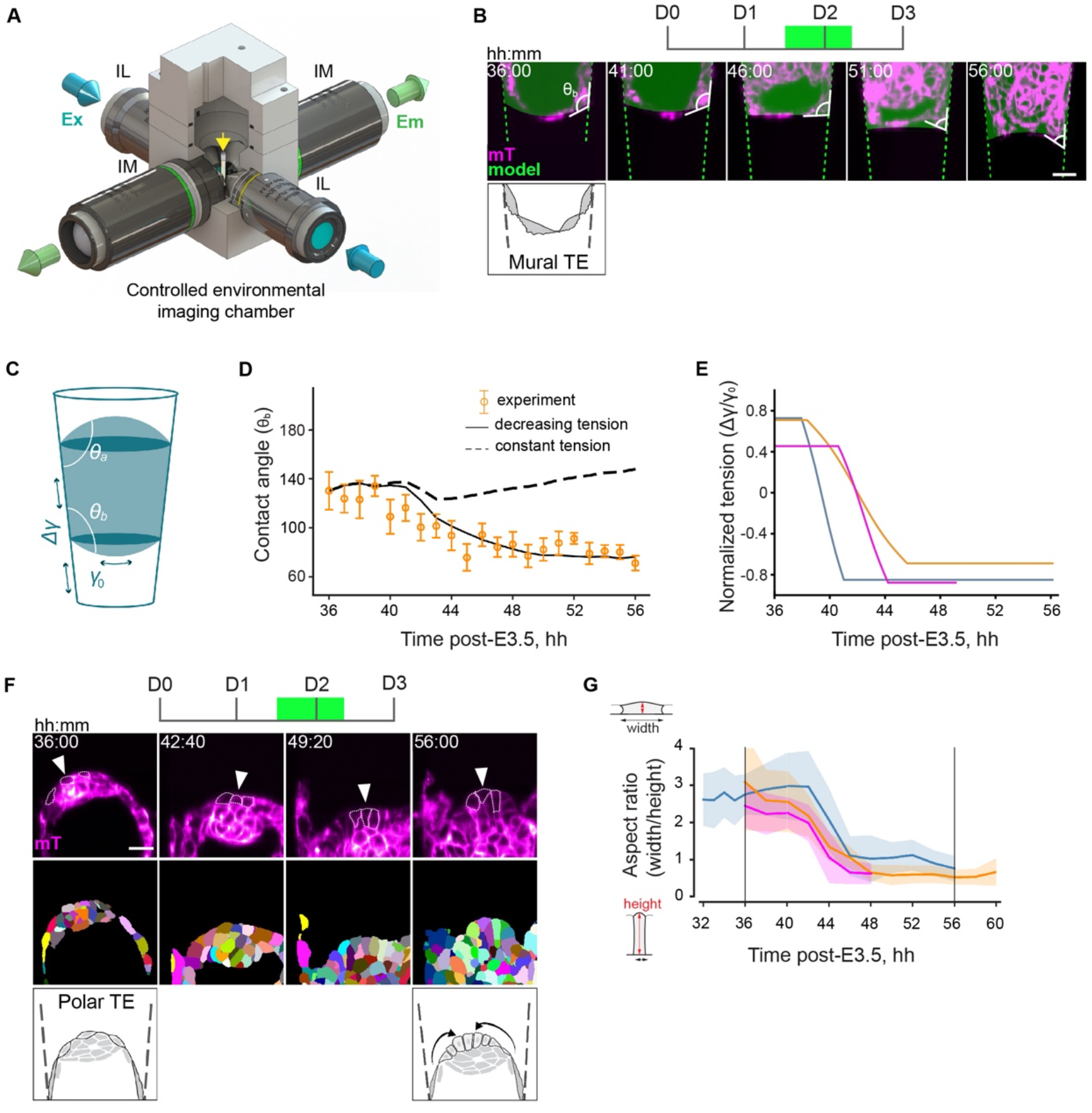
A droplet-wetting process can explain embryo-uterus interactions upon implantation. **(A)** Schematic of the MuVi-SPIM setup with two low-NA illumination objective lenses (IL), two high-NA imaging objective lenses (IM), and the controlled environmental imaging chamber with the sample holder (yellow arrow). **(B)** Representative time-lapse images of the mural TE of the mTmG developing embryo; mTomato (magenta). Fitted droplet model (embryo) and the frustum shape (crypt) shown in green; contact angle (θ) between mural TE/culture medium and mural TE/crypt interfaces is shown. **(C)** Schematic of the active droplet in a frustum-shaped confinement. The tension difference Δ_γ_ between substrate-droplet and substrate-medium interfaces and the droplet-medium tension γ_0_ are shown as green arrows; θ_a_ and θ_b_ denote top and bottom contact angles, respectively. **(D)** Simulated contact angle dynamics for constant tension (dashed line) and decreasing tension (solid line) with experimental data points (orange points). **(E)** Dynamics of the normalized embryo-substrate interfacial tension difference. Colors correspond to independent experiments. **(F)** Top, representative single-plane time-lapse images of the polar TE of mTmG developing embryo; mTomato (magenta). Exemplar pTE cells are marked with arrowheads, cell perimeter is outlined. Middle, corresponding images of the 3D cell membrane segmentation. Bottom, the schematic of pTE cell columnarization and invagination. **(G)** Dynamics of the width-to-height aspect ratio of the pTE cells. Colors correspond to independent experiments. Average values across 15-20 cells per time point (solid line) and standard deviations (shaded area) are shown. Scale bars, 20 μm. t =00:00, hours: minutes from recovery at E3.5. See also Materials and Methods, Supplemental Note, Figure S4, Video S7

We therefore modelled the embryo as a fluid droplet, confined within a conical frustum representing the 3E-uterus. The droplet has different interfacial tensions with the substrate and the medium, with γ_0_ denoting the tension of the droplet-medium interface and Δγ the tension difference between droplet-substrate and substrate-medium interfaces, and it is subject to a Laplace pressure ΔP that acts as a Lagrange multiplier to the imposed droplet volume V. This model predicts a relationship between the interfacial tensions, contact points, and contact angles between the droplet (embryo) and the substrate (uterus), given the volume of the droplet (Figure 4B-C, S4A-E, Supplemental Note).

We measured the volume from live imaging data of the embryo, and calculated the contact-angle dynamics in the presence and absence of adhesion changes between the droplet and the substrate using simulation-based inference (Tejero-Cantero et al., 2020) to determine the remaining parameter values of our model (Figure S4E-F, Table S3). Experimental measurement of the contact angle between the embryo and the 3E-uterus surface showed a remarkable agreement with the theoretical values for increasing adhesion (Figure 4D, Figure S4G, H; Video S7). This suggests that the tissue-scale shape dynamics resulting from embryo implantation can be biophysically understood as an active wetting process. This model further predicts that failure to adhere to the uterus should lead to maximum contact angles or near-spherical embryo shapes, in line with the outcome of embryo culture in the hydrogel without RGD modification (Figure 2A, Figure S4D).

Notably, the postulated tension release at the embryo-substrate interface exactly corresponds to the condition required for pTE cells to constrict apically, invaginate and form ExE – which had been achieved *ex vivo* only by removing the mTE and releasing tension acting on the pTE (Bedzhov et al., 2014; Ichikawa et al., 2022). Light-sheet microscopy and measurement of the pTE cell aspect ratio showed that pTE cells indeed undergo apical constriction (Figure 4F-G) within the predicted time interval (Figure 4E), indicating that the 3E-uterus recapitulates the embryo-uterus interaction, releasing the TE tension and enabling the development of the whole embryo *ex vivo* for the first time.

Collectively, these findings show that the embryo-uterus tissue-level interaction upon implantation can be biophysically described as a droplet-wetting process and that this embryo-uterus interaction releases tension acting on the TE, enabling ExE formation.

### Multi-view light-sheet microscopy reveals peri-implantation egg cylinder growth dynamics

To dissect the coordination between embryo growth, TB migration, and the uterus, we further characterized the growth dynamics of the tissues comprising the egg cylinder, using the MuVi-SPIM. As a result of TE tension release as described above, CDX2-GFP;mTmG embryos underwent ExE invagination and proliferation as well as egg cylinder patterning (Figure 5A, B). Live imaging of H2B-GFP; mTmG embryos showed tissue growth at cellular resolution without compromising egg cylinder proliferation and patterning (Figure 5C, Figure S5A, B) (Christodoulou et al., 2018; Ichikawa et al., 2022). The egg cylinder elongated along the M/AM axis at a rate of 5.52 μm/h with its tip moving at 4.62 μm/h (Figure 5C, D, Video S9). EPI cell lineage tracks estimated an average cell-cycle length of 8:38 hh:min (Figure 5E), and EPI tissue volume increased 1.78x over 8 hours on average (Figure 5F). Thus, our engineering approach of hydrogel microfabrication combined with multi-view light-sheet imaging revealed the cellular dynamics of embryonic and extraembryonic tissues and allowed us to quantitively characterize their substantial growth and dynamic morphogenesis upon embryo-uterus interaction.

### Collective trophoblast migration delineates uterine space for embryo development

Collectively, these findings are in agreement with the model in which adhesion of the embryo to the uterine matrix induces a wetting process at the tissue scale, enabling TE tension release, and collective migration of TB cells at the cellular scale, displacing Reichert’s membrane and generating a space for the egg cylinder growth. This suggests an intricate interaction and coordination between embryos and the uterine tissues at cell and tissue scales. Since the 3E-uterus *ex vivo* system not only enables us to quantitatively characterize the cellular and tissue dynamics, as described above, but also to perturb the system with spatio-temporal control, we tested our coordination model by experimental manipulation of its key parameters, the TB migration velocity and geometry of the uterus.

First, the TB migration velocity should be fast enough for the displacement of Reichert’s membrane to accommodate egg cylinder tissue growth and morphogenesis. On the contrary, the model predicts that if the TB migration velocity is slower, egg cylinder tissue growth and morphogenesis will be physically disrupted by Reichert’s membrane. As TB cells undergo collective migration, we hypothesized that their migration may be dependent on Rac1 (Rac family small GTPase 1), that is also required for collective cell migration during mouse anterior-posterior axis specification (Migeotte et al., 2010) and gastrulation (Migeotte et al., 2011; Saykali et al., 2019). Genetic perturbation of Rac1^-/-^ reportedly shows growth retardation at E5.75 and arrest during gastrulation (Migeotte et al., 2010; Sugihara et al., 1998). In accordance with it, 3E-uterus culture revealed that elongation of the embryo is significantly limited in Rac1^-/-^ embryos (Figure 6A, B). Notably, the growth of EPI tissue is compromised, and its elongation appears to be blocked by Reichert’s membrane (Figure 6A, C). Pharmacological inhibition of Rac1 by NSC23766 (Gao et al., 2004) showed retention of the TB migration front in a reversible manner, compared to the control embryos (Figure S6A, B). Together, these results indicate that TB migration is dependent on Rac1 and is required for EPI tissue growth, in agreement with the model prediction.

**Figure 5.**
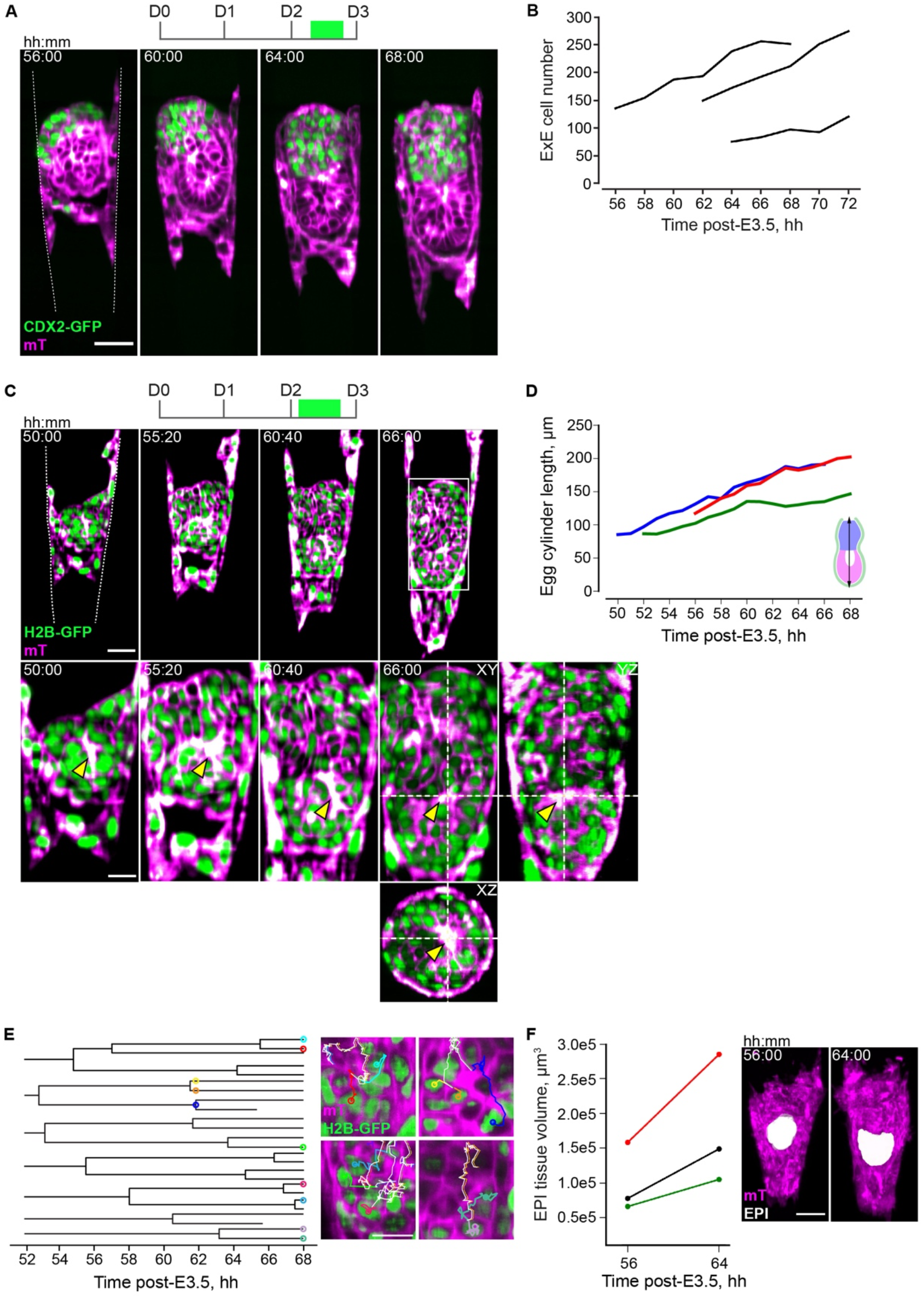
Multi-view light-sheet microscopy reveals peri-implantation egg cylinder growth dynamics. **(A)** Representative single-plane time-lapse images of the CDX2-GFP;mTmG embryo developing in 3E-uterus; CDX2-GFP (green), mTomato (magenta). See also Video S8. **(B)** Quantification of the ExE cell numbers. **(C)** Representative single-plane time-lapse images of the H2B-GFP;mTmG embryo developing in 3E-uterus. Bottom, 2x zoom into the epiblast region. Right, YZ and XZ image sections, showing 3D resolution. GFP (green), mTomato (magenta). The crypt surface is outlined. The yellow arrow marks rosettes leading to formation of the pro-amniotic cavity. See also Video S9. **(D)** Change in egg cylinder length between 50 h and 68 h after recovery at E3.5. Colors correspond to independent experiments. **(E)** Epiblast cell lineage dendrograms. Right, corresponding cells marked as dots with different colors overlaying the dendrograms and the image slices; cell lineage tracks are depicted as a 2D overlay. (F) Change in epiblast tissue volume between 56 h and 64 h after recovery at E3.5. Colors correspond to independent experiments. Right, 3D image of the developing embryo with segmented EPI tissue volume; mTomato (magenta), EPI tissue segmentation (white). Scale bars, 50 μm, 25 μm (C, bottom), 20 μm (E, right). t =00:00, hours: minutes after recovery at E3.5.

Next, the displacement velocity of Reichert’s membrane depends on the geometry of the 3E-uterus, too. To interfere with it, we placed the embryo upside-down in the crypt (Figure 6D-F). Due to the crypt diameter gradient, the model predicts that this would slow down the velocity of Richert’s membrane displacement even if the TB migration velocity remains unchanged, leading to impairment of the egg cylinder growth. Measurement of the displacement of the embryonic pole and Reichert’s membrane front revealed that egg cylinder elongation is accompanied by a parallel Reichert’s membrane dislocation in 3E-uterus at the rate of 4.2 μm/h (Figure 6E). However, in the upside-down orientation, the diplacement of Reichert’s membrane relative to the egg cylinder growth is slower (Figure 6F) and the egg cylinder growth appeared to be eventually blocked (Figure 6G). These data indicate that the spatial coordination is disrupted by a change in the 3E-uterus geometry, leading to the impairment of the EPI morphogenesis in agreement with the model prediction.

**Figure 6.**
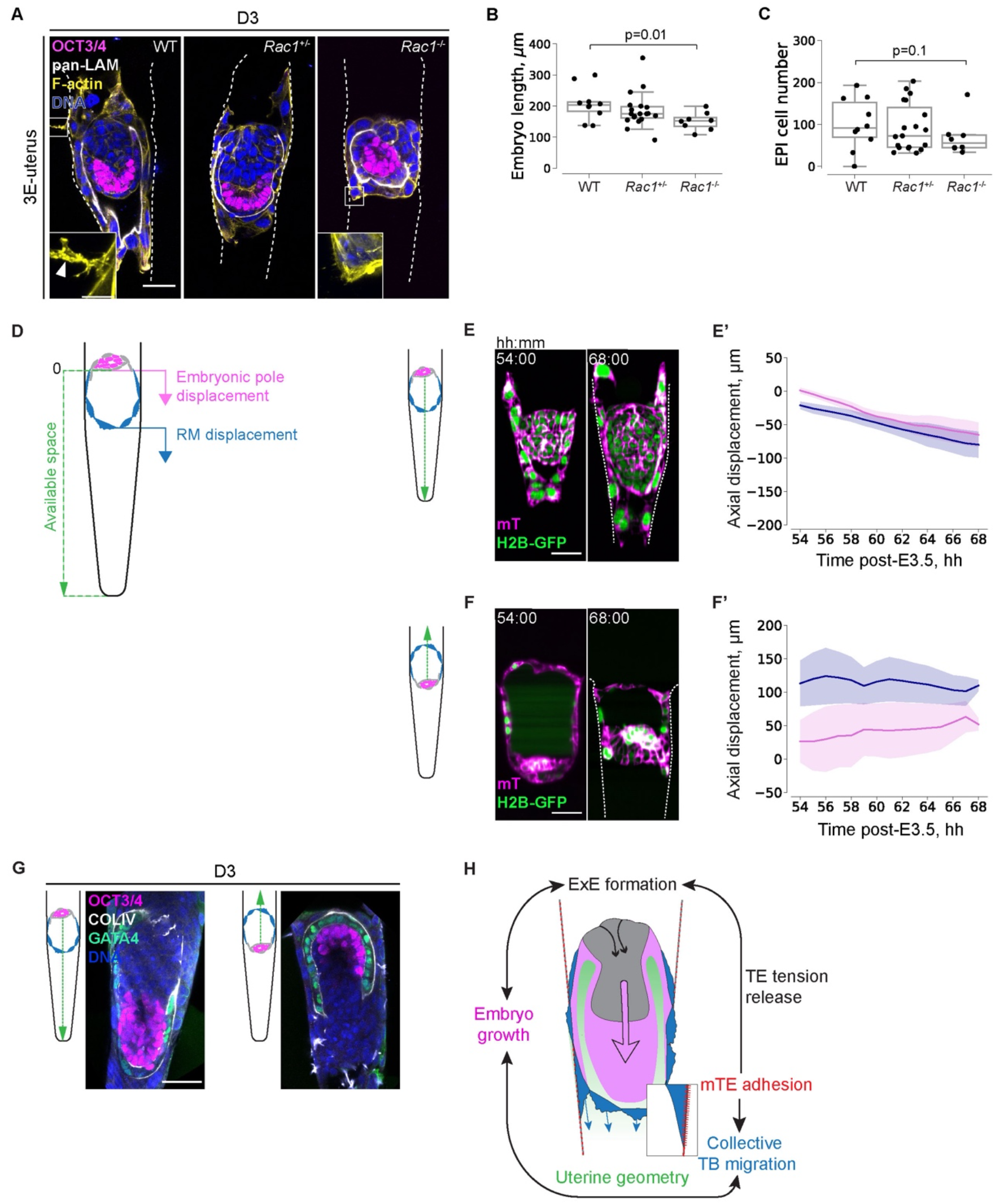
Collective trophoblast migration delineates uterine space for embryo development. **(A)** Left to right, representative immunofluorescence images of WT, Rac1^+/-^, and Rac1^-/-^ embryos, cultured up to day 3 (D3) in 3E-uterus. Simultaneous staining for OCT3/4 (magenta), pan-Laminin (pan-LAM, white), F-actin (yellow), and nuclei (DNA, blue). The crypt surface is outlined. **(B)** Boxplots showing embryo length in WT, Rac1^+/-^, and Rac1^-/-^ embryos. n = 10 (WT), 19 (Rac1^+/-^), 8 (Rac1^-/-^), all embryos pooled from three experimental replicates. Data points correspond to embryos, midline marks the median, boxes indicate interquartile range. P-values were calculated using Mann-Whitney U test. **(C)** Boxplots showing epiblast cell number between WT, Rac1^+/-^, and Rac1^-/-^ embryos. n = 10 (WT), 19 (Rac1^+/-^), 8 (Rac1^-/-^), all embryos pooled from three experimental replicates. Data points correspond to embryos, midline marks the median, boxes indicate interquartile range. P-values were calculated using Mann-Whitney U test. **(D)** Schematic of the quantified parameters, embryonic pole displacement (pink), RM displacement (blue) along the crypt axis (green). Coordinates are scaled to the starting coordinate of the embryonic pole. **(E)** Representative time-lapse images of H2B-GFP;mTmG embryo developing in a downward orientation; GFP (green), mTomato (magenta). **(E’)** Average displacement of the embryonic pole (pink line) and RM (blue line) along the crypt axis in the downward developing embryo. Shaded regions indicate standard deviation. **(F)** Representative time-lapse images of H2B-GFP;mTmG embryo developing in an upward orientation; GFP (green), mTomato (magenta). **(F’)** Average displacement of the embryonic pole (pink line) and RM (blue line) along the crypt axis in the upward developing embryo. Shaded regions indicate standard deviation. **(G)** Representative immunofluorescence images of 3E-uterus embryos from day 3 in the downward (left) and upward (right) orientations, simultaneously stained for OCT3/4 (magenta), GATA4 (green), and nuclei (DAPI, blue). **(H)** The proposed model for the coordination underlying peri-implantation mouse embryo development. mTE adhesion to the uterine tissue triggers TE tension release, resulting in ExE formation, and collective mTE/TB migration which delineates uterine space for embryo growth and morphogenesis. Scale bars, 50 μm. t =00:00, hours: minutes from recovery at E3.5.

Together, these experimental perturbations support the model in which the coordination between the embryo growth, TB collective migration and the uterine geometry plays a key role in periimplantation mouse development (Figure 6H).

## DISCUSSION

In this study, we developed a new engineered uterus, 3E-uterus, which allowed us for the first time to culture the whole peri-implantation mouse embryo *ex vivo,* recapitulating the embryo-uterus interaction and supporting the differentiation of TB cells and Reichert’s membrane. Combined with light-sheet microscopy, this system allows for monitoring the cellular dynamics and perturbing cellular process by genetic, pharmacological, and biophysical means, in order to gain insights into the underlying mechanisms. Our study revealed that integrin-mediated adhesion of TB cells onto the uterine matrix not only releases TE tension to drive its invagination and ExE formation, but also induces collective migration of TB cells. This TB cell migration, in turn, displaces Reichert’s membrane so that the embryo has space for growth and morphogenesis. These findings thus reveal a dynamic coordination between embryo growth, TB cell migration and uterine geometry that plays an essential role in peri-implantation mouse development (Figure 6H).

Our finding that the embryo adheres to and interacts with the uterus for its morphogenesis is reminiscent of recent studies demonstrating a key role of attachment of the blastoderm or endoderm to the vitelline envelope in insect morphogenesis (Bailles et al., 2019; Münster et al., 2019). In the implanting mouse embryo, however, adhesion of the TB cells induces loss of polarity, possibly epithelial-mesenchymal transition (Damjanov et al., 1986; Lamouille et al., 2014), and collective migration.

To study the mechanics of these embro-uterus interactions, we used theoretical concepts from the physics of wetting. Theoretical approaches for classical wetting as well as active tissue wetting have yielded key insights into different multicellular spreading phenomena (Douezan et al. 2011, Gonzales-Rodriguez et al. 2012, Alert and Casademunt 2019, Perez-Gonzales et al. 2019). Here we modeled the embryo as a confined simple droplet with adaptive adhesion to the substrate, and inferred the substrate-adhesion dynamics from the observed shape changes (Figure 4C-F, Figure S4, Supplemental Note). Our results suggest that embryo implantation occurs through a biologically tuned capillarity-like process (Figure 4E, Figure S4H, Video S7).

Together, our findings revealed an intricate interaction and coordination between embryo development and uterine dynamics upon mammalian implantation both in space and time. The space for proper embryo growth is defined by the dynamic interaction between the extra-embryonic and uterine tissues, whereas the timing of adhesion adaptation matches invagination and generatino of extra-embronic ectoderm, which in turn influences embryonic growth (Ichikawa et al. 2022). These results open up new avenues for exploring dynamics of the tissue-tissue and embryo-extraembryonic-tissue interactions in many other contexts.

When studying complex tissue-tissue interactions, one approach is to use engineered systems to recapitulate certain aspects of the tissue-tissue interactions. The advantage of this method is to be able to control the components of the system, hence providing insights into its mechanisms. Our recent study showed that tissue properties and geometry guide patterning and morphogenesis of intestinal organoids via robust spatial gradients of the cell density and signaling (Gjorevski et al., 2022). In this study, a similar microfabrication recapitulated the uterine environment, with the possiblity to control its geometry, stiffness, and adhesion properties, and revealed the mechanism of how the integrin-mediated adhesion and tissue geometry coordinate peri-implantation embryonic development.

The engineering of the uterus can be developed further. More bio-degradable and dynamic hydrogels may help accommodate embryo development to more advanced stages (Brassard and Lutolf, 2019; Chrisnandy et al., 2022; Qazi et al., 2022). The uterine endometrium certainly provides a complex composition of mechanical and biochemical signaling (Dey et al., 2004), and incorporating more cellular components into the engineering platform may improve the *ex vivo* culture and our understanding of the mechanisms of embryo-uterus interactions.

The interaction between embryos and the uterus *in utero,* however, likely involves dynamic changes in the uterine tissue too (Kelleher et al., 2018; Flores et al., 2020; Ueda et al., 2020). This cannot be recapitulated by the present 3E-uterus, and may require more complete reconstruction of the uterine environment (Boretto et al., 2017; Turco et al., 2017). Alternatively, or ultimately, studying the physiological embryo-uterus interaction needs direct monitoring of the process *in utero* (Huang et al., 2020). These new methods will open an exciting perspective for studying the feto-maternal interaction, which is key for improving reproductive medicine, as well as gaining insights into its co-adaptation over the evolutionary time-scale.

## SUPPLEMENTAL FIGURES

**Figure S1.**
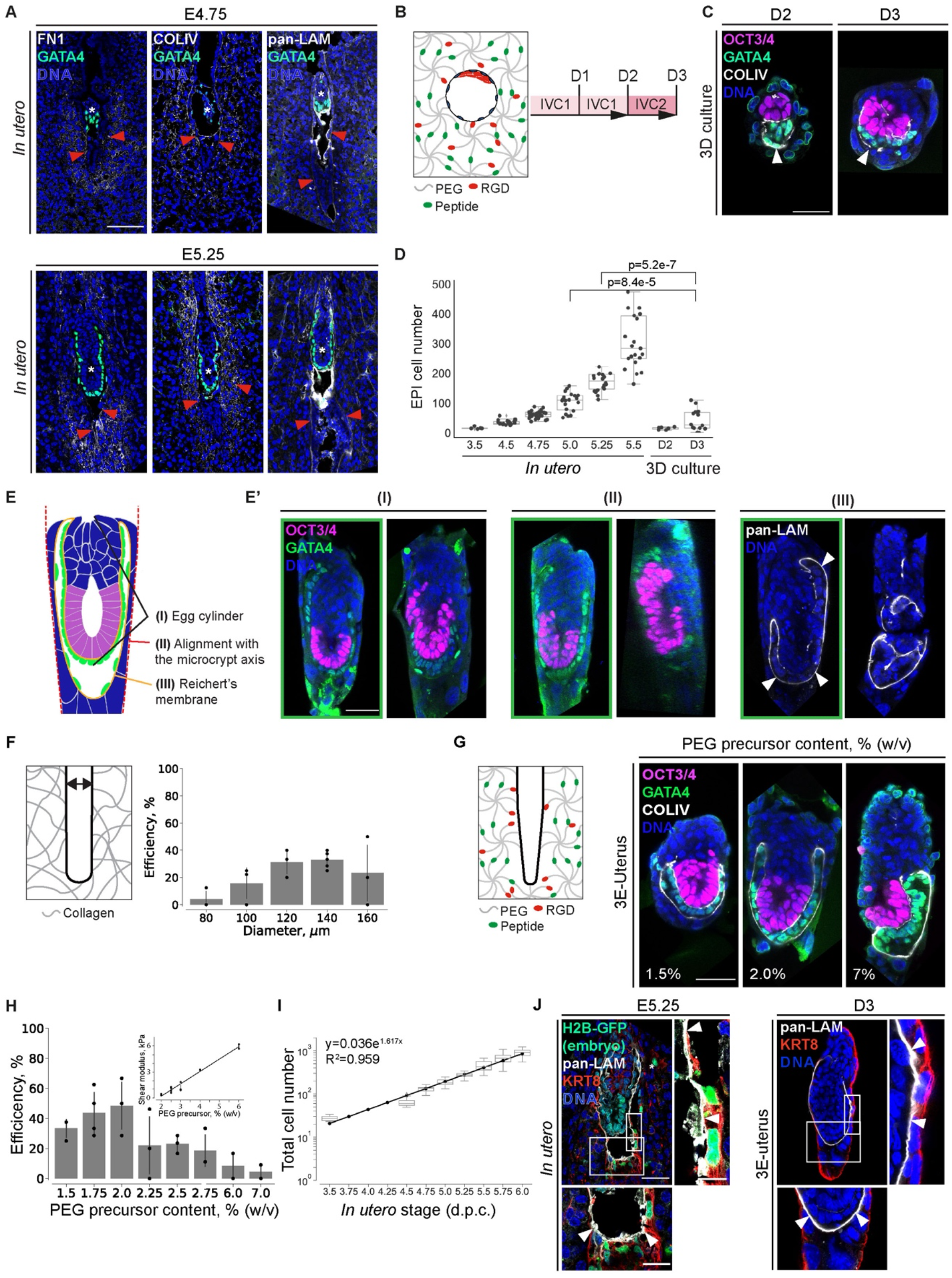
*Ex vivo* Engineering Uterine Environment with topographically patterned hydrogels. **Related to Figure 1** **(A)** Immunofluorescence images of the representative E4.75 (top) and E5.25 (bottom) uteri cross-sections, stained for Fibronectin (FN1, white), Collagen IV (COLIV, white), and Laminin (LAM, white) (from left to right), GATA4 (green), and nuclei (DNA, blue). White asterisks mark implanted embryos; sectioning thickness, 20 μm. Red arrowheads point at the uterine basal membrane. **(B)** Schematic of the experimental design. Embryos, recovered at E3.5 and treated with Tyrode’s solution to remove Zona pellucida (ZP), are embedded for 3D culture into the drops of hydrogel on the day of recovery (D0), and cultured for up to three days (D1-3). The hydrogel is comprised of RGDSPG (Arg-Gly-Asp-Ser-Pro-Gly, red)-functionalized 8-arm Poly(ethylene glycol) (PEG, gray), crosslinked via the metalloprotease-cleavable peptides (Peptide, green). IVC1 and ICV2 stand for “In Vitro Culture” medium 1 and 2, respectively. Inner cell mass (ICM), red; trophectoderm (TE), blue. **(C)** Representative immunofluorescence images of the embryos embedded and cultured 3D inside hydrogel drops until day 2 (D2) and day 3 (D3), stained for OCT3/4 (magenta), GATA4 (green), and nuclei (DNA, blue). White arrowheads point at Reichert’s membrane. **(D)** Comparison of the epiblast (EPI) cell numbers between the embryos developed *in utero* until E3.5 – E5.5 (Ichikawa et al., 2022) and the embryos embedded and cultured 3D inside hydrogel drops until days 2-3 (D2-3). n = 7 (D2) and n = 14 (D3). P-values are calculated using Mann-Whitney U test. **(E)** Schematic of the embryo morphology criteria (I-III), based on which the efficiency of the *ex vivo* culture is evaluated. **(E’)** Representative immunofluorescence images of the embryos after 3E-uterus culture for 3 days. Immunostaining for OCT3/4 (magenta), GATA4 (green), Laminin (LAM, white), and nuclei (DNA, blue). The embryos that form egg cylinder (I), show the egg cylinder axis in line with the crypt axis (II), and form Reichert’s membrane (III), are considered to be successfully developed (outlined in green; 46%; n = 12 of 26, pooled from three experimental replicates). White arrowheads point at Reichert’s membrane, **(F)** Left, the scheme of diameter evaluation using cylindrical crypts inside Collagen I hydrogel. Right, barplots showing average 3E-uterus efficiency for culture inside cylindrical crypts with different diameters. n = 2 (80 μm), n = 3 (100 μm), n = 3 (120 μm), n = 5 (140 μm), n = 3 (160 μm), indicates numbers of experimental replicates. Whiskers mark standard deviations. **(G)** Left, the scheme of the funnel-shape crypt inside PEG hydrogel with a diameter gradient. Right, representative immunofluorescence images of the 3E-uterus embryos from day 3 (D3) grown in 1.5%, 2%, 2.5%, and 7% PEG precursor concentrations (from left to right), stained for OCT3/4 (magenta), GATA4 (green), Colagen IV (COLIV, white), and nuclei (DNA, blue). **(H)** Barplots showing 3E-uterus efficiency at day 3 at various PEG precursor contents. n = 3 (1.5%), n = 4 (1.75%), n = 3 (2%), n = 3 (2.25%), n = 3 (2.5%), n = 3 (2.75%), n = 3 (6%), n = 2 (7%), indicates a number of experimental replicates. Whiskers mark standard deviations. Inset, rheological measurement showing linear relationship between the PEG precursor content (%, w/v) and the Shear modulus (kPa). **(I)** Plot showing total cell number (EPI+VE) against *in utero* developmental stage defined by the time of recovery. The days of 3E-uterus culture were matched with the *in utero* stages based on the log-linear regression. Equation of the regression line for total cell number (EPI and VE) is y = 0.133e1.489x; that for EPI cell number is y = 0.036e1.617x. n = 6 (E3.5), n = 21 (E4.5), n = 28 (E4.75), n = 20 (E5.0), n = 20 (E5.25), n = 21 (E5.5), n = 21 (E5.75) and 22 (E6.0), data from (Ichikawa et al., 2022). Y scale, log 10. **(J)** Left, representative immunofluorescence image of E5.25 uterus cross-section from the F1 female mated with the H2B-GFP male. Immunostaining for Cytokeratin 8 (KRT8, red), pan-Laminin (pan-LAM, white), GFP (green), and nuclei (DNA, blue). White asterisk marks TB cell, distant from the Reichert’s membrane. Right, 4x zoom; bottom, 2x zoom. Right, representative immunofluorescence image of the 3E-uterus embryo from day 3 (D3). Immunostaining for Cytokeratin 8 (KRT8, red), pan-Laminin (pan-LAM, white), and nuclei (DNA, blue). Right, 4x zoom; bottom, 2x zoom. White arrowheads point at Reichert’s membrane. Scale bars, 50 μm, 25 μm (2x zoom), 12.5 μm (4x zoom).

**Figure S2.**
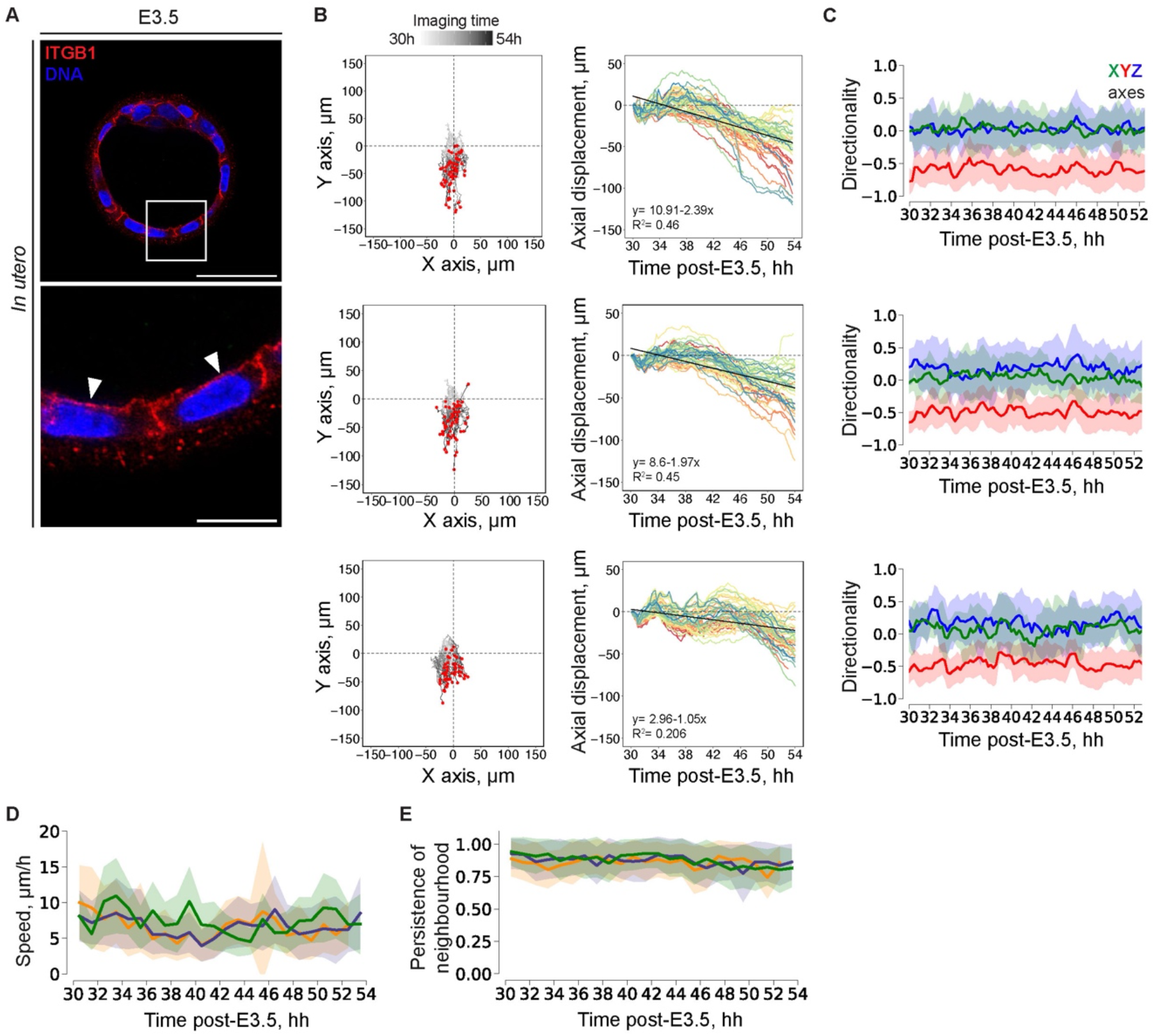
Integrin localization in the blastocyst and collective trophoblast cell migration. **(A)** Representative immunofluorescence image of the blastocyst-stage embryo (E3.5), simultaneously stained for integrin beta 1 (ITGB1, red), and nuclei (DAPI, blue). Bottom, 4x zoom. Arrowheads point to the basal integrin localization in TE. **(B)** Left, mural TE cell trajectories for three different embryos; coordinates in XY plane are normalized to the starting coordinates. End coordinates are marked with red dots. Right, displacement of mural TE cells along the Y-axis vs imaging time post-E3.5. From top to the bottom, n = 61, 58, 51, respectively. The linear regression fit is shown as a black line. **(C)** Directionality of the mTE/TB migration along the X, Y, and Z axes (green, red, and blue, respectively) between subsequent time points (every hour of live imaging) for three embryos (from top to bottom). n = 61,58, 51, respectively. **(D)** TB migration speed (μm/h) vs imaging time post E3.5. Colors correspond to the three embryos from B and C, **(E)** Persistence of the nearest mTE/TB four-cell neighbourhood between subsequent time points (every hour of live imaging).

**Figure S3.**
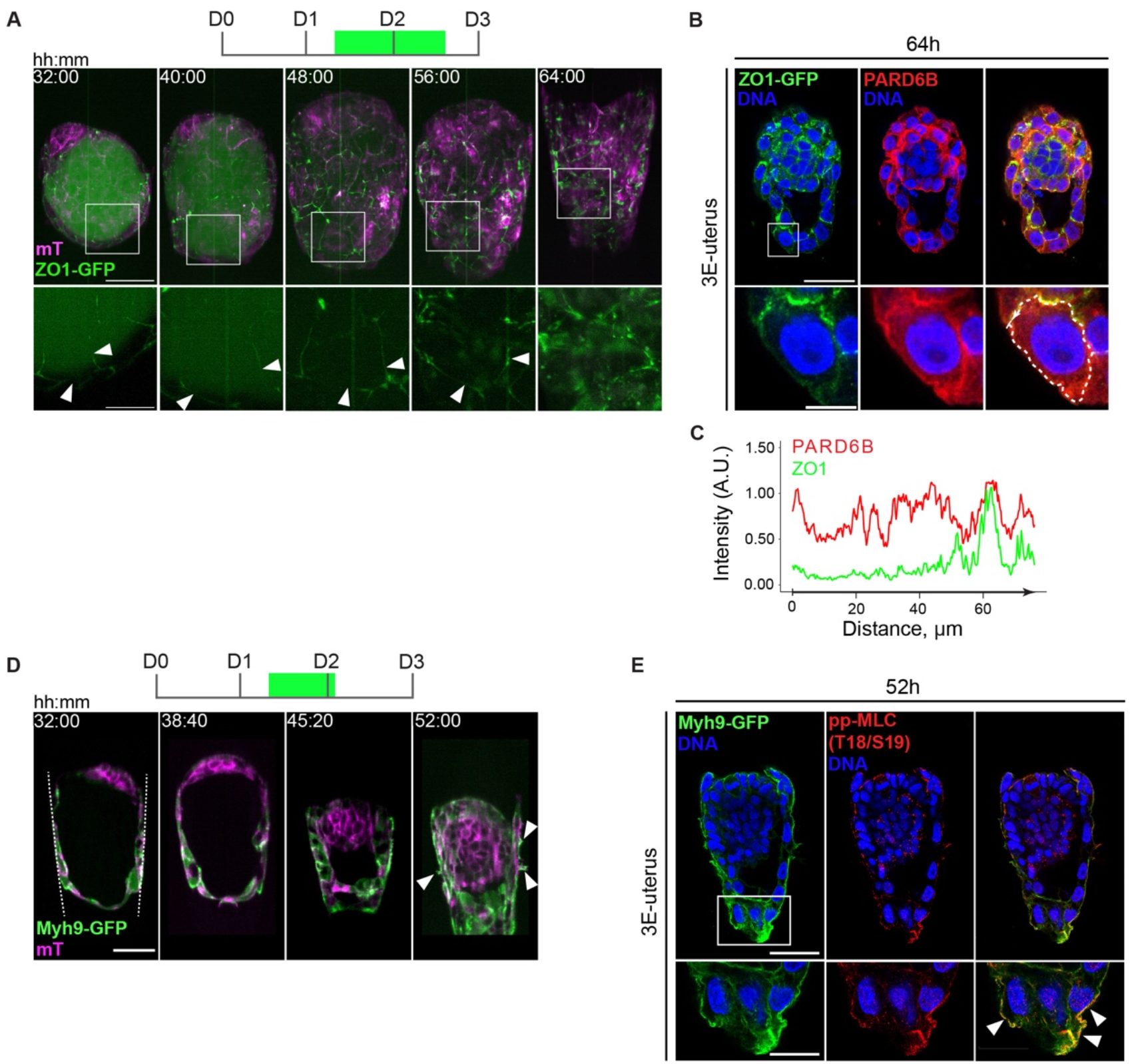
Trophoblast cells acquire mesenchymal motility upon adhesion. **(A)** Representative 3D projections of time-lapse images of ZO1-GFP;mTmG developing embryo; ZO1-GFP (green), mTomato (magenta). Bottom, 2.5x zoom into a TE cell; white arrowheads mark cell-cell interface. See also Video S3. **(B)** Representative immunofluorescence images of the 3E-uterus embryo after live imaging, simultaneously stained for ZO1-GFP (green) PARD6B (red), and nuclei (DNA, blue). From left to right, ZO1-GFP, PARD6B, composite image channels. Bottom, 4x zoom of the TB cell. **(C)** Intensity profile of ZO1 and PARD6B signals along the cell surface outlined in B (bottom), including apical and basolateral regions. **(D)** Representative single-plane time-lapse images of the Myh9-GFP;mTmG developing embryo; Myh9-GFP (green), mTomato (magenta). The crypt surface is outlined. See also Video S6. **(E)** Representative immunofluorescence images of the 3E-uterus embryo after live imaging, simultaneously stained for Myh9-GFP (green) phospho-MLC (T18/S19) (red), and nuclei (DNA, blue). From left to right, Myh9-GFP, phospho-MLC (T18/S19), composite image channels. Bottom, 2x zoom. White arrowheads point at the apical TB cell surface. Scale bars, 50 μm, 20 μm (2.5x zoom), 12.5 μm (4x zoom). t=00:00, hours: minutes from recovery at E3.5.

**Figure S4.**
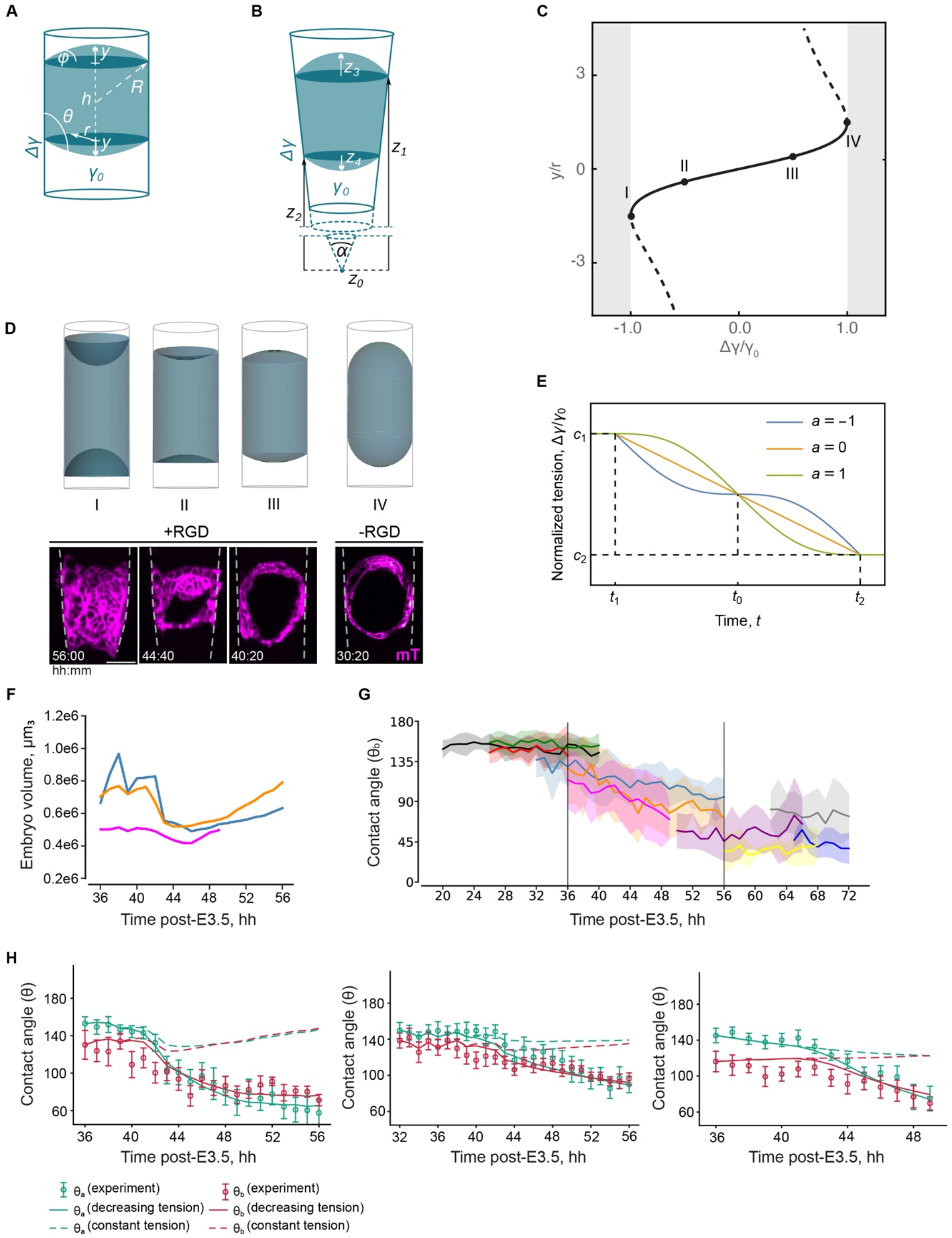
Characterization of the droplet-wetting model. **(A-B)** Geometry of the droplet model in cylindrical and conical confinement. **(A)** The equilibrium shape of the droplet in a cylindrical confinement of radius *r* is described by the distance *h* between the two contact lines and by the height *y* of the top and bottom spherical caps. These caps can also be characterized by the curvature radius *R* and angle *φ*. The contact angle *θ* depends on the droplet-medium tension *γ*_0_ and the Young tension Δγ. **(B)** The droplet in a conical frustum with angle *α* is described by the positions of the top and bottom contact lines *z*_1_, *z*_2_ measured from the conical tip *z*_0_ = 0, and by the heights of the top and bottom spherical caps *z*_3_ and *z*_4_, respectively. When the caps curve into the embryo, their heights assume negative values. **(C)** Bifurcation diagram for the equilibrium solutions Eq. S6 (Supplemental Note) of the droplet in cylindrical confinement. The solid line corresponds to the stable solution *y-,* whereas the dashed line denotes the unstable branch *y*_+_. **(D)** Calculated equilibrium shapes of the droplet in cylindrical confinement (top) at the transition to total wetting (I), in the regime of partial wetting (II, III), and dewetting (IV) at points shown in (C). Example single-plane time lapse images (bottom) from control conditions show embryo shape dynamics indicating increasing adhesion, whereas embryos in non-adherent conditions (without RGD in the hydrogel) recapitulate shapes predicted for the dewetting condition. Scale bars, 50 μm. **(E)** Sigmoid model of the Young tension adaptation Eq. S16 drawn for three values of the modulation amplitude *a* ∈ [-1, 1]. Constants *c*_1_ and *c*_2_ specify the initial and final values of the normalized tension. The adaptation begins at a time instance *t*_1_ and ends at a time instance *t*_2_. A full specification of the model requires five constants, e.g. the mid time *t*_0_ = (*t*_1_ + *t*_2_) / 2, the duration *Δt* = *t*_2_ – *t*_1_, the constants *c*_1_ and *c*_2_, and the amplitude *a*. **(F)** Volume dynamics in the developing embryos between 36 h and 56 h after E3.5. Colors correspond to different embryos; n = 3. **(G)** Contact angle (θb) dynamics in developing embryos. Colors correspond to different embryos imaged in time intervals between 20 h and 72 h after E3.5; n = 10. **(H)** From left to right, simulated contact angle dynamics for constant tension (dashed line), and decreasing tension (solid line), with experimental data points (green and red points for θa and θb, respectively) for three different embryos between 36 h and 56 h from recovery at E3.5.

**Figure S5.**
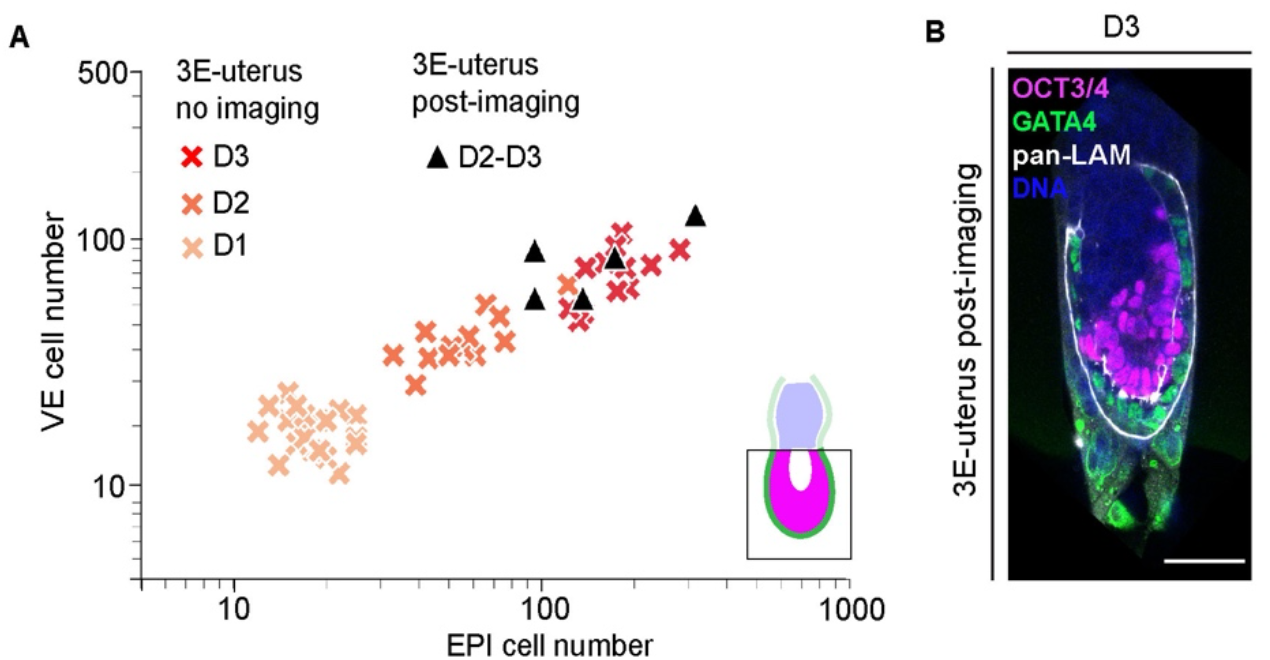
Evaluation of embryo morphology after live imaging. **(A)** Scatterplot showing the numbers of epiblast (EPI) cells (x-axis) vs the numbers of visceral endoderm (VE) cells (y-axis) that cover EPI in the 3E-uterus embryos developed in the incubator for three days (D1-D3, no imaging), and 3E-uterus embryos developed in the incubator, but then live imaged with MuVi-SPIM for 20 – 24 hours up to day 3 (D3, after imaging). N = 12, pooled from three experimental replicates (D3, no imaging), n = 5, pooled from five experimental replicates (D3, after imaging). The groups of imaged and not imaged D3 embryos did not significantly differ in terms of EPI (P = 0.69) and VE (P=0.37) cell numbers. P-values were calculated using t-test. XY scale, log 10. **(B)** Representative immunofluorescence image of the day 3 embryo after live imaging with MuVi-SPIM. Staining for OCT3/4 (magenta), GATA4 (green), pan-Laminin (pan-LAM, white), and nuclei (DNA, blue). Scale bar, 50 μm.

**Figure S6.**
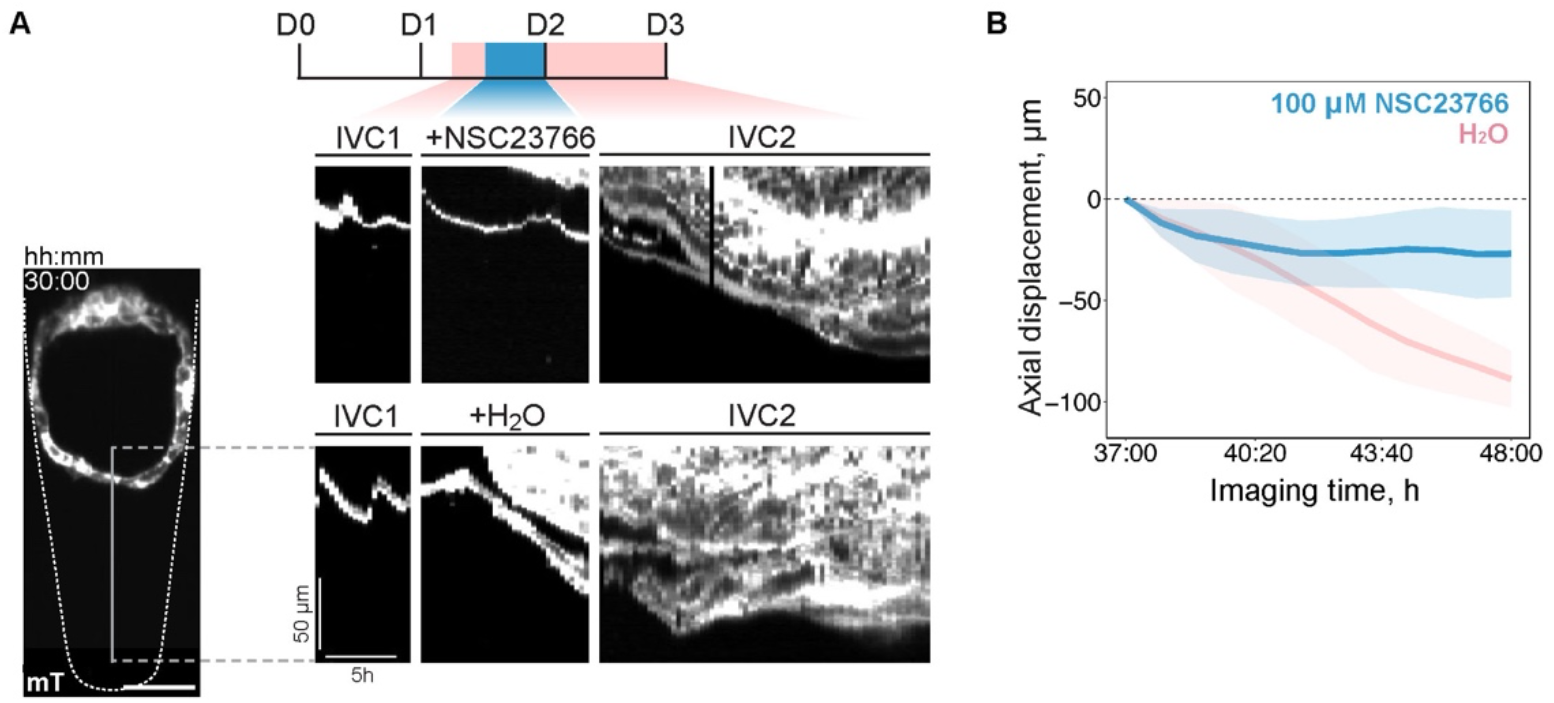
Pharmacological inhibition of TB migration. **(A)** Kymographs showing mural TE leading-edge displacement along the Y-axis, indicated with the solid line on the left-most panel. Embryos from the same litter were incubated with 100 μM NSC23766 (top) and water (bottom) in IVC1 between 37 h and 48 h after recovery at E3.5; mTomato (grey). **(B)** Mural TE leading-edge displacement along the Y-axis in embryos, incubated with 100 μM NSC23766 (blue) and water (pink) in IVC1 between 37 h and 48 h. n = 4, 4, respectively. Average values (solid lines) and standard deviations (shaded area) are shown.

## SUPPLEMENTAL VIDEOS

**Video S1. *<u>E</u>x vivo* <u>e</u>ngineering uterine <u>e</u>nvironment with biomimetic hydrogel topography (3E-uterus) for peri-implantation mouse embryo culture. Related to Figure 1.**

00:05 – 00:15 (mm:ss), engineered crypt dimensions and crypt microfabrication; 0:15 – 0:25, embryo embedding and *ex vivo* culture. 0:32 – 0:50, 3E-uterus recapitulates peri-implantation egg cylinder morphology. 3D projections and the corresponding single-plane immunofluorescence images of the embryo developed *in utero* until E5.25 (left), and the embryo developed *ex vivo* until day 3 (right). Simultaneous staining for OCT3/4 (magenta), GATA4 (green), Collagen IV (COLIV, white), and nuclei (DNA, blue). 0:50 – 0:55, 3E-uterus recapitulates differentiation of trophoblast (TB). Left, single-plane immunofluorescence image of the E5.25 uterus cross-section. Right, single-plane immunofluorescence image of the 3E-uterus embryo from day 3 (D3). Simultaneous immunostaining for TFAP2C (yellow) and nuclei (DNA, blue). Yellow arrows point at the TB cells. Scale bars, 25 μm.

**Video S2. mTE/TB cells migrate collectively upon adhesion. Related to Figure 2.**

3D projection of the time-lapse video of H2B-GFP in the embryo developing in the crypt made of PEG with RGD; H2B-GFP (green to yellow). Trajectories of individual mTE cells are marked with red lines. t =00:00, hours: minutes after recovery at E3.5. Scale bar, 50 μm.

**Video S3. Tight junctions rearrange in the migratory mTE/TB. Related to Figure S3.**

3D projection of the time-lapse video of developing ZO1-GFP;mTmG embryo; ZO1-GFP (green), mTomato (magenta). t =00:00, hours: minutes after recovery at E3.5. Scale bar, 50 μm.

**Video S4. Migratory mTE/TB exhibits apical actin-rich protrusions. Related to Figure 3.**

Time-lapse video of the developing Lifeact-GFP;mTmG embryo; Lifeact-GFP (green), mTomato (magenta). t=00:00, hours: minutes after recovery at E3.5. Scale bar, 50 μm.

**Video S5. mTE forms dynamic lamellipodia. Related to Figure 3.**

3D projection of the time-lapse video of Lifeact-GFP in the mTE of the developing embryo; Lifeact-GFP (spectral). t=00:00, minutes:seconds after recovery at E3.5. Scale bar, 20 μm.

**Video S6. Nonmuscle myosin 9 dynamics in the migratory mTE/TB. Related to Figure S3.**

Time-lapse video of the developing Myh9-GFP;mTmG embryo; Myh9-GFP (green), mTomato (magenta). t=00:00, hours: minutes after recovery at E3.5. Scale bar, 50 μm.

**Video S7. Fit of the droplet-wetting model with the experimental data. Related to Figure 4.**

Top, from left to right, time-lapse videos of the three developing mTmG embryos; mTomato (magenta). t=00:00, hours: minutes after recovery at E3.5. Scale bar, 50 μm. Bottom, corresponding droplet-wetting models of the embryos in conical frustum shapes.

**Video S8. Multi-view light-sheet live microscopy reveals ExE invagination. Related to Figure 5.**

Time-lapse video of the developing CDX2-GFP;mTmG embryo; CDX2-GFP (green), mTomato (magenta). t=00:00, hours: minutes after recovery at E3.5. Scale bar, 50 μm.

**Video S9. Cellular dynamics and growth of the whole peri-implantation embryo. Related to Figure 5.**

Time-lapse video of the developing H2B-GFP;mTmG embryo; H2B-GFP (green), mTomato (magenta). t=00:00, hours: minutes after recovery at E3.5. Scale bar, 50 μm.

## SUPPLEMENTAL TABLES

**Table S1.**
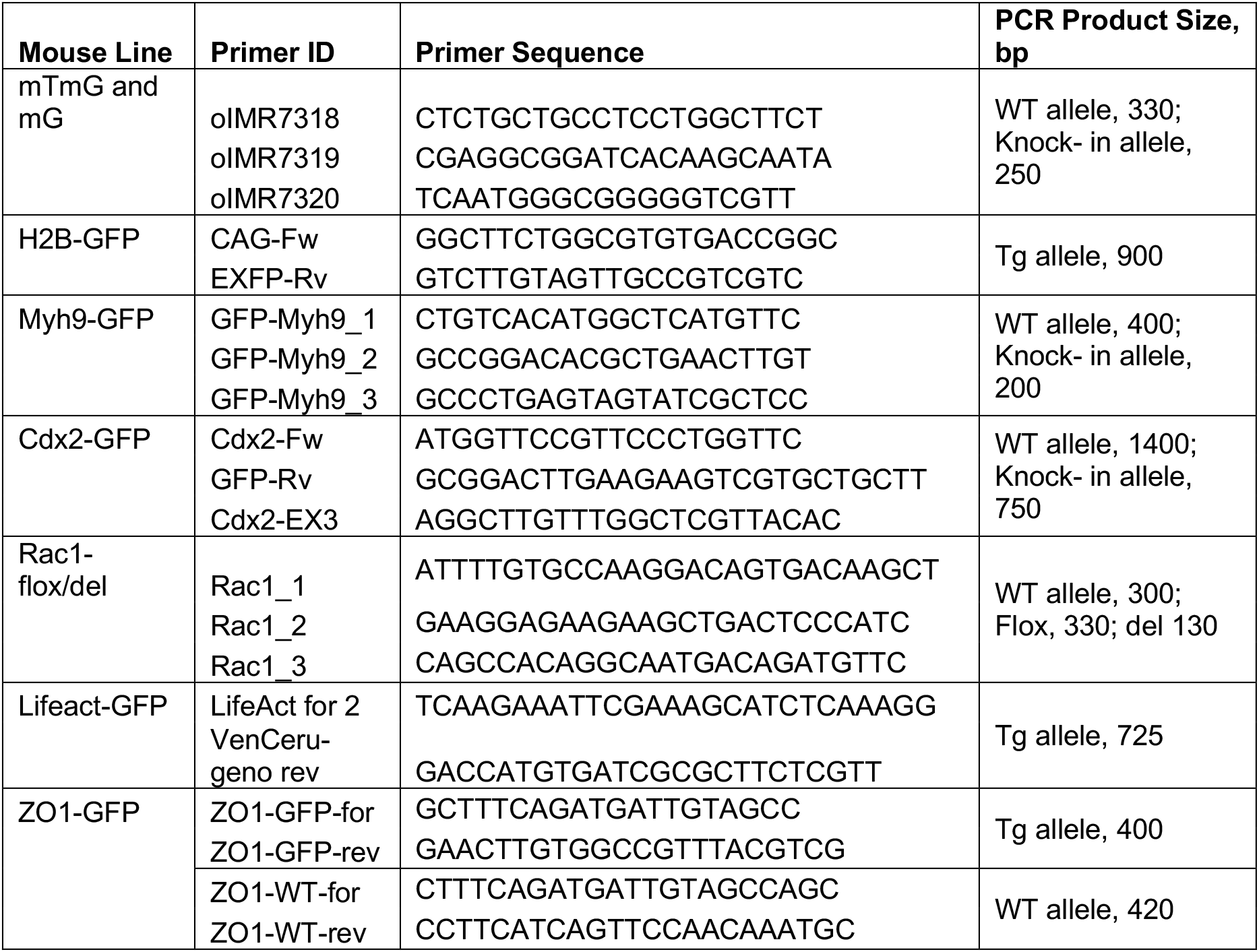
Genotyping primers and PCR product sizes.

**Table S2.** Summary of the microscopy types and imaging settings.

| Figure | Panel | Microscopy | Lasers, nm | Voxel size, $\mu\text{m}$ (XYZ) |
| --- | --- | --- | --- | --- |
| **1** | A | Confocal, airyscan | 405, 488, 633 | 0.0824 x 0.0824 x 0.1917 |
|  | B | Confocal, airyscan | 405, 488, 633 | 0.0824 x 0.0824 x 0.1917 |
|  | D | Confocal | 405, 488, 546, 633 | 0.207 x 0.207 x 1 |
|  | E | Confocal | 405, 488, 546, 633 | 0.207 x 0.207 x 1 |
|  | I | Confocal | 405, 546 | 0.207 x 0.207 x 1 |
|  | J | Confocal | 405, 546 | 0.207 x 0.207 x 1 |
| **2** | A | Confocal | 405, 546, 633 | 0.232 x 0.232 x 1 |
|  | B | Confocal | 405, 546, 633 | 0.232 x 0.232 x 1 |
|  | C | Confocal | 405, 488, 633 | 0.232 x 0.232 x 1 |
|  | F | InVi-SPIM | 488, 561 | 0.104 x 0.104 x 1.000 |
|  | F' | InVi-SPIM | 488, 561 | 0.104 x 0.104 x 1.000 |
| **3** | A | Confocal, airyscan | 405, 488, 633 | 0.0824 x 0.0824 x 0.1917 |
|  | C | Confocal, airyscan | 405, 488, 633 | 0.0824 x 0.0824 x 0.1917 |
|  | E | InVi-SPIM | 488, 561 | 0.104 x 0.104 x 1.000 |
|  | E' | InVi-SPIM | 488, 561 | 0.104 x 0.104 x 1.000 |
|  | F | Confocal, airyscan | 405, 488 | 0.0824 x 0.0824 x 0.1917 |
| **4** | C | MuVi-SPIM | 561 | 0.295 x 0.295 x 1.000 |
|  | G | MuVi-SPIM | 561 | 0.295 x 0.295 x 1.000 |
| **5** | A | MuVi-SPIM | 488, 561 | 0.295 x 0.295 x 2.000, 1.000 |
|  | C | MuVi-SPIM | 488, 561 | 0.295 x 0.295 x 1.000 |
|  | E | MuVi-SPIM | 488, 561 | 0.295 x 0.295 x 1.000 |
|  | F | MuVi-SPIM | 488, 561 | 0.295 x 0.295 x 1.000 |
| **6** | A | Confocal | 405, 488, 546, 633 | 0.6919 x 0.6919 x 2 |
|  | E | MuVi-SPIM | 488, 561 | 0.295 x 0.295 x 2.000, 1.000 |
|  | F | MuVi-SPIM | 488, 561 | 0.295 x 0.295 x 2.000, 1.000 |
|  | G | Confocal | 405, 488, 546, 633 | 0.207 x 0.207 x 1 |
| **S1** | A | Confocal, airyscan | 405, 488, 633 | 0.0824 x 0.0824 x 0.1917 |
|  | C | Confocal | 405, 488, 546, 633 | 0.1389 x 0.1389 x 1 |
|  | E' | Confocal | 405, 488, 546, 633 | 0.207 x 0.207 x 1 |
|  | G | Confocal | 405, 488, 546, 633 | 0.207 x 0.207 x 1 |
|  | J | Confocal, airyscan | 405, 488, 546, 633 | 0.0824 x 0.0824 x 0.1917 |
| **S2** | A | Confocal, airyscan | 405, 633 | 0.0824 x 0.0824 x 0.1917 |
| **S3** | A | InVi-SPIM | 488, 561 | 0.104 x 0.104 x 1.000 |
|  | B | Confocal | 405, 488, 633 | 0.207 x 0.207 x 1 |
|  | D | InVi-SPIM | 488, 561 | 0.104 x 0.104 x 1.000 |
|  | E | Confocal, airyscan | 405, 488, 633 | 0.0824 x 0.0824 x 0.1917 |
| **S4** | D | MuVi-SPIM | 561 | 0.295 x 0.295 x 2.000, 1.000 |
| **S5** | B | Confocal | 405, 488, 546, 633 | 0.207 x 0.207 x 1 |
| **S6** | A | InVi-SPIM | 561 | 0.208x0.208x1.000 |

**Table S3.**
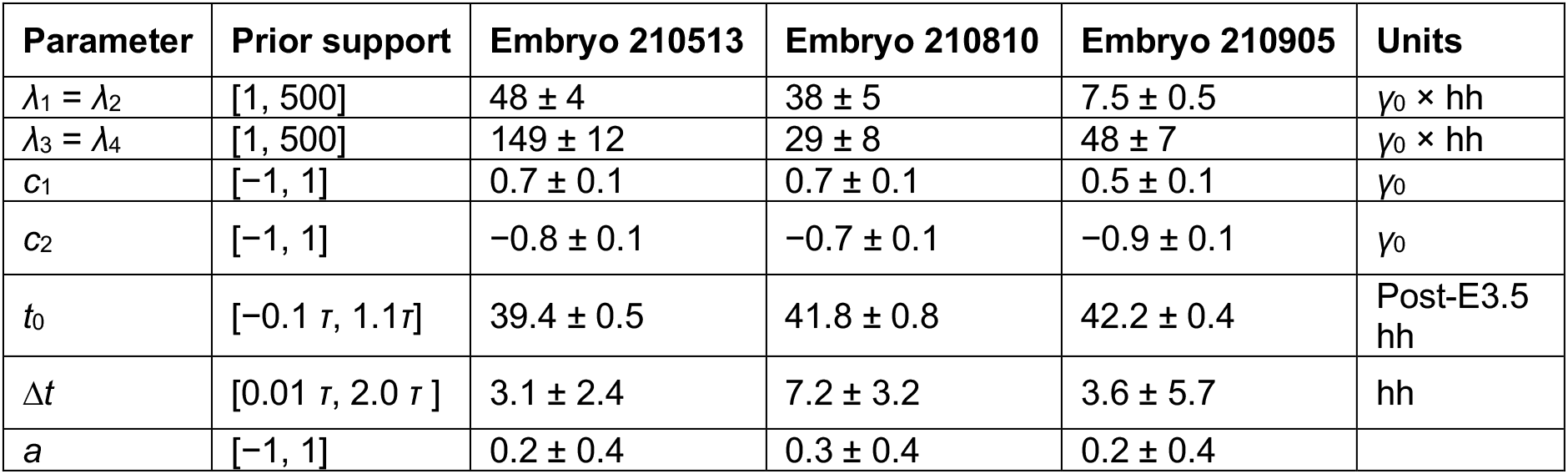
Simulation-based inference of the model parameters values: dissipative coefficients for the positions of contact lines λ_1,2_ and for the heights of embryo caps λ_3,4_; the initial and final values of the Young tension c_1_ and c_2_ respectively; mid time t_0_, duration Δ_t_ and modulation parameter a of the time-dependent Young tension Δγ [Supplemental Note, Eq. (S16)]. Values of to have been converted to the post-E3.5 time. Error bounds are given by one standard deviation.

| Parameter | Prior support | Embryo 210513 | Embryo 210810 | Embryo 210905 | Units |
| --- | --- | --- | --- | --- | --- |
| $\lambda_1 = \lambda_2$ | [1, 500] | $48 \pm 4$ | $38 \pm 5$ | $7.5 \pm 0.5$ | $\gamma_0 \times \text{hh}$ |
| $\lambda_3 = \lambda_4$ | [1, 500] | $149 \pm 12$ | $29 \pm 8$ | $48 \pm 7$ | $\gamma_0 \times \text{hh}$ |
| $c_1$ | [-1, 1] | $0.7 \pm 0.1$ | $0.7 \pm 0.1$ | $0.5 \pm 0.1$ | $\gamma_0$ |
| $c_2$ | [-1, 1] | $-0.8 \pm 0.1$ | $-0.7 \pm 0.1$ | $-0.9 \pm 0.1$ | $\gamma_0$ |
| $t_0$ | $[-0.1 \tau, 1.1 \tau]$ | $39.4 \pm 0.5$ | $41.8 \pm 0.8$ | $42.2 \pm 0.4$ | Post-E3.5<br>hh |
| $\Delta t$ | $[0.01 \tau, 2.0 \tau]$ | $3.1 \pm 2.4$ | $7.2 \pm 3.2$ | $3.6 \pm 5.7$ | hh |
| $a$ | [-1, 1] | $0.2 \pm 0.4$ | $0.3 \pm 0.4$ | $0.2 \pm 0.4$ | |

## MATERIALS AND METHODS

### Animal Work

All animal work was performed in the Laboratory Animal Resources (LAR) at the European Molecular Biology Laboratory (EMBL) with permission from the Institutional Animal Care and Use Committee (IACUC) overseeing the operation (IACUC number TH11 00 11). LAR is operated according to the Federation of European Laboratory Animal Science Associations (FELASA) guidelines and recommendations. All mice were housed in IVC cages in pathogen-free conditions with 12-12 hours light-dark cycle and used for experiments at the age of 8 to 35 weeks.

### Mouse Lines and Genotyping

The following mouse lines were used in this study: a F1 hybrid strain between C57BL/6 and C3H (B6C3F1) as wild-type (WT), Cdx2-GFP (Mcdole and Zheng, 2012), mTmG (Muzumdar et al., 2007), H2B-GFP (Hadjantonakis and Papaioannou, 2004), Lifeact-GFP (Riedl et al., 2010), GFP-Myh9 (Zhang et al., 2012), ZO1-GFP (Foote et al., 2013). Rac1^flox/flox^ conditional allele (Walmsley et al., 2003) was crossed with ZP3-Cre line (Lewandoski et al., 1997) to generate Rac1^+/-^ animals. To obtain zygotic Rac1^-/-^ embryos, Rac1^+/-^ females were crossed with Rac1^+/-^ males. Standard tail genotyping procedures were used to genotype transgenic mice (see Table S1 for primers and PCR product sizes).

### Mouse Embryos

Female estrous cycle synchronization was used to increase the natural mating efficiency (Whitten M. K., 1957). The embryonic day 0.5 (E0.5) was defined as noon on the day when a vaginal plug was detected. Pre-implantation mouse embryos were flushed from the uteri of the plugged females with 37 °C KSOMaa with HEPES (Zenith Biotech, ZEHP-050, 50 ml) using a syringe equipped with a cannula (Acufirm, 1400 LL 23). Embryos were handled using an aspirator tube (Sigma, A5177), connected to a glass pipette pulled from a glass microliter pipette (Blaubrand intraMark 708744). Procedures were performed under a stereomicroscope (Zeiss, StreREO Discovery.V8) equipped with a thermal plate (Tokai Hit) at 37°C (Behringer et al., 2014).

Peri-implantation embryos were dissected from uteri in DMEM (Gibco, 11880028) supplemented with 15% heat-inactivated FBS (PAA, A15-080), 2 mM GlutaMAX (Gibco, 35050061), and 10 mM HEPES (Sigma, H0887), as described (Nagy et al., 2003).

### LDTM Hydrogel Precursor Synthesis

Low-defect thiol-Michael addition (LDTM) PEG hydrogel was synthesized and characterized according to the previously published study (Rezakhani et al., 2020). Briefly, to synthesize peptide-functionalized PEG macromers (PEG-PEP), vinyl sulfone-functionalized 8-arm PEG (8-arm PEG-VS) and the peptide Ac-GCRE-GPQGIWGQ-ERCG-NH2 (mol wt 1773.97 g/mol) with matrix metalloproteinases sensitive sequence (GPQGIWGQ) were dissolved in triethanolamine (TEA) (0.3 M, pH 8.0), and the 8-arm PEG-VS was added dropwise to the excess of peptides (VS/SH=10) and reacted for two hours at room temperature under inert atmosphere. The reaction solution was dialyzed (Snake Skin, molecular weight cut-off 10K) against ultrapure water (pH < 7) for 5 days at 4°C, and the final product was lyophilized. The lyophilized product was dissolved in water to make 10% precursor solutions.

### Hydrogel Formation

LDTM hydrogels were formed by Michael type addition of PEG-PEP precursors onto 8-arm PEG-VS. To make hydrogel networks of desired final PEG content, proper volumes of 10% (w/v) 8-arm PEG-VS in TEA and 10% (w/v) PEG-PEP in water were mixed in molar stoichiometric ratio of VS/SH=0.8. For example, to make 100 μL of LDTM hydrogels of 2.5% (w/v), taking into account each precursors’ densities, 8.8 μL of 8-arm PEG-VS, 10 μL of TEA buffer, 65 μL of distilled water and 16.20 μL of PEG-PEP were mixed. For conditions containing RGD adhesion peptide (Ac-GRCGRGDSPG-NH2, mol wt 1002.04 g/mol), different volumes of RGD were added to the mix before addition of the PEG-PEP precursor, and the molar ratio of VS/TH was adjusted as VS/(TH-RGD)=0.8. The table below shows the mixing values for LDTM gels (2.5% (v/w)) with different RGD contents.

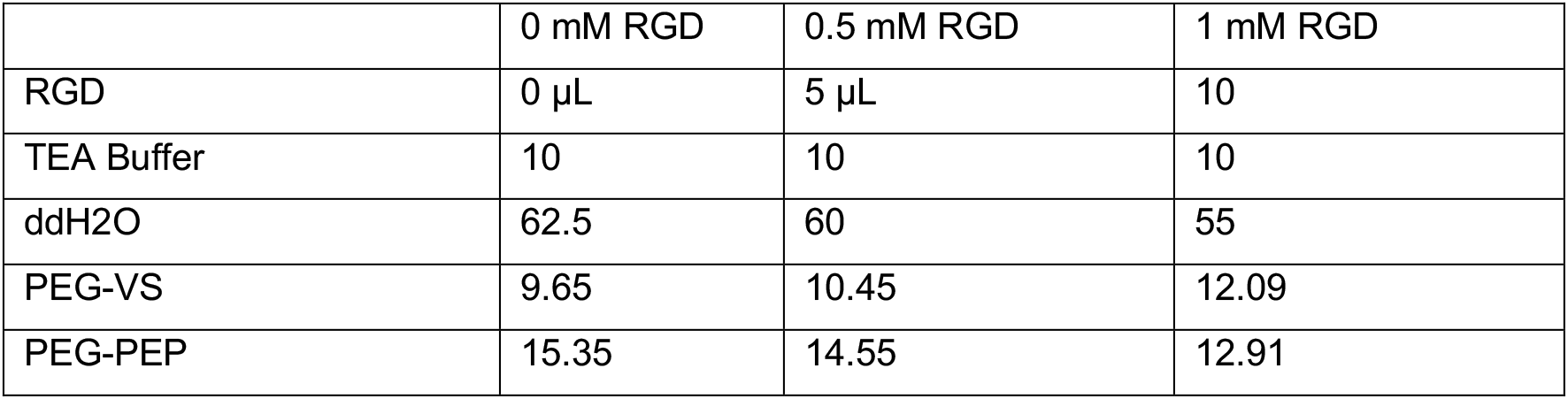

### Rheological Measurements of Hydrogels

The shear modulus (G’) of hydrogels was determined by performing small-strain oscillatory shear measurements on a Bohlin CVO 120 rheometer with plate-plate geometry. Briefly, 1–1.4 mm thick hydrogel discs were prepared and allowed to swell in water overnight. The mechanical response of the hydrogels sandwiched between the parallel plates of the rheometer was recorded by performing frequency sweep (0.1–10 Hz) measurements in a constant strain (0.05) mode at room temperature.

### Hydrogel Holders and Microtopography Stamps Fabrication

For hydrogel casting on the bottom of the 3.5 cm dish, special ring-like PDMS holders were fabricated (see “Fabrication of topographically patterned hydrogels”). These hydrogel holders defined resulting hydrogel thickness to 1mm and had overhanging features to hold the hydrogel block attached to the bottom of the plate. Holders were fabricated using conventional soft-lithography methods established at the Center of Micronanotechnology (CMi, EPFL). In brief, the array of circular rings (∅ = 10 mm) was drawn using a CleWin (Phoenix Software). The designed layout was written with a diode laser onto a fused silica plate coated with chrome and positive photoresist (Nanofilm) using an automated system (VPG200, Heidelberg Instruments). Exposed photoresist was removed with a developer (DV10, Süss MicroTec) and the chrome layer underneath was etched with an acid–oxidizer solution of perchloric acid, cerium ammonium nitrate and water. The resulting mask was developed with TechniStrip P1316 (Microchemicals) to remove the residual resist and extensively washed with ultra-pure water. The mold was made from double-layered epoxy-based negative photoresist SU8. First, a 250 μm thick layer of SU8 GM1075 (Gerlteltec) photoresist was cast onto a dehydrated silicon wafer using a negative resist coater (LMS200, Sawatec). After baking at 110 °C for 4 hours, a second 250 μm thick layer of the SU8 GM1075 was coated, resulting in total thickness of a 500 μm. After second bake, this wafer was aligned and exposed to ultraviolet (UV) radiation through the mask (MA6/BA6, Süss MicroTec). After the postexposure bake at 85 °C for 5 hours, the wafer was developed with propylene glycol monomethyl ether acetate (Sigma) and baked at 135 °C for 4 h. The wafer was then plasma-activated and silanized with vapored trichloro (1H,1H,2H,2H-perfluorooctyl) silane (Sigma-Aldrich) overnight. This wafer was then used for polydimethylsiloxane (PDMS) moulding (Sylgard 184, Dow Corning). Ten weight-parts of elastomer base were vigorously mixed with one part of curing agent and poured onto the mould. After degassing under vacuum, PDMS was baked for 24 h at 80 °C. The resulting PDMS replica was cut and punched with appropriate size biopsy punchers (5 mm for inner hydrogel area, 18 mm outside diameter). Resulting hydrogel holders were sterilized with UV and kept sterile until further use.

The stamps featuring micro-pillars topography were fabricated using conventional soft-lithography methods established at the Center of Micronanotechnology (CMi, EPFL). In brief, the 3D models of the micropillars were designed in Autodesk Inventor. The STL model was further processed in DeScribe 1.7 (Photonic Professional) to optimize it for printing 2PP lithography (NanoScribe GT2, Photonic Professional). The printing parameters were defined to have an slicing distance of 1 um and a hatching distance of 600 nm. Writing was performed in the galvo scan mode and the 3D model was divided in 400 x 400 μm sub-blocks, corresponded to the FOV of 25X objective. The model was printed in acrylic photopolymer resin IP-S (Photonic Professional), which was deposited on the surface of indium-tin oxide (ITO)-coated soda lime slides. Once the printing process was finalized, the slides were developed for 7 min in propylene glycol monomethyl ether acetate (PGMEA, Sigma-Aldrich), followed by 5 min rinsing with ultrapure isopropanol and gentle blow-drying. Then samples were UV cured for 10 min (MA6/BA6, Süss MicroTec) and backed in the 80 °C oven for 3 hours. To fabricate a master mold featuring inverted topography, 3D printed models were then plasma-activated and silanized with vapored trichloro (1H,1H,2H,2H-perfluorooctyl) silane (Sigma-Aldrich) overnight. Then polydimethylsiloxane (PDMS, Sylgard 184, Dow Corning) was used for moulding. Ten weight-parts of elastomer base were vigorously mixed with one part of curing agent and poured onto the mould. After degassing under vacuum, PDMS was baked for 12 h at 80 °C. The resulting PDMS replica was cut, plasma-activated and silanized with vapored trichloro (1H,1H,2H,2H-perfluorooctyl) silane (Sigma-Aldrich) overnight. This replica was used multiple times as a master for molding stamps, following the same protocol (PDMS Sylgard 184, 1:10 ratio, baked at 80 °C for 12 h).

### Fabrication of Topographically Patterned Hydrogels

Elastomeric stamps containing the desired geometries in bas-relief were coated with bovine serum albumin in PBS (1% w/v in PBS; Thermo Fisher Scientific) for overnight to prevent hydrogel adhesion. Before use stamps were washed once with distilled water and dried with the gentle air blowing. For hydrogel casting, PDMS ring holders were placed on the bottom of the 3.5 cm dish and UV sterilized prior use. Then a drop of liquid hydrogel precursor (see “Hydrogel Formation”) was made in the center of the ring spacer and then a stamp with microtopography was placed atop. After 30-40 min polymerization in the incubator (37 C) stamps were removed and hydrogels were covered with PBS. Hydrogels were used either the same day or stored for about one week.

### Embryo Culture

#### 3E-uterus

In Vitro Culture medium (IVC1 and IVC2) was prepared as described (Bedzhov et al., 2014). Hydrogels with microfabricated crypts (see “Hydrogel Formation” and “Fabrication of topographically patterned hydrogels”) were equilibrated in 3 ml of IVC1 medium in an incubator with a humidified atmosphere of 5% CO_2_ at 37 °C (Thermo Scientific, Heracell 240i) for at least 12 h prior to 3E-uterus embryo culture. After recovery at noon on embryonic day 3.5 (E3.5), embryos were serially transferred to IVC1 microdrops in a culture dish covered with mineral oil (Sigma, M8410-1L). The time of embryo culture was counted from the time of embryo recovery (D0 = E3.5). In approximately one hour after recovery, embryos were briefly treated in Tyrode’s solution (Sigma, T1788) to remove *Zona pellucida,* washed repeatedly (Behringer et al., 2014), and left in a culture dish with IVC1 medium for at least an hour inside the incubator. Zona-free embryos were carefully positioned inside microfabricated hydrogel crypts in a downward mTE orientation with a fused tip of a thin glass pipette. The medium was exchanged to IVC1 in 24 h (Day 1) and to IVC2 in 48h (Day 2).

#### 3D hydrogel-embedded culture

LDTM PEG hydrogel components were mixed on ice. 15 μl of the mix was added to an inner well of a pre-warmed μ-Slide Angiogenesis dish, and embryos were carefully transferred and mixed inside the hydrogel (2-3 embryos per drop). To prevent embryos from adhering to glass or reaching gel surface, the dish was flipped regularly during gel solidification inside the incubator. 35 μL of pre-warmed IVC1 medium were then added to each well. Subsequently, the medium was exchanged, and then exchanged again to IVC1 in 24 h (Day 1) and to IVC2 in 48h (Day 2).

### Single Embryo Genotyping

Individual embryos were mouth pipetted into 200 μl PCR tubes containing 10 μl of lysis solution of 200 μg/ml Proteinase K in Taq polymerase buffer (Thermo Fischer Scientific, B38). The lysis reaction was carried out for 1 h at 55°C, followed by 10 min at 96°C. The resulting genomic DNA was mixed with relevant primers (Table S1) for determination of genotype via PCR.

### Immunofluorescence Preparation and Staining

Recovered embryos were fixed with 4% paraformaldehyde (Electron microscopy sciences, 19208) in PBS for 15 minutes at room temperature. For *ex vivo* cultured embryos, the hydrogel with embryos was gently dissected and fixed with 4% paraformaldehyde for 30 minutes at room temperature with agitation. For immunostaining of active integrin and di-phosphorylated myosin regulatory light chain (ppMRLC), fixation was performed in 1% PFA in PBS supplemented with MgCl_2_. The samples were subsequently washed in PBST buffer (0.1% Tween-20 in PBS; Sigma, 85113), *ex vivo* cultured embryos were carefully dissected from the hydrogel at this step. Permeabilization was performed with 0.5% Triton X-100 (Sigma, T8787) in PBS for 30 minutes at room temperature with gentle agitation. After several washes in the wash buffer (2.5% BSA (Sigma, A9647) in PBST), embryos were incubated in the blocking buffer (5% BSA in PBST) overnight at 4°C. Embryos were stained with primary antibodies diluted in the blocking buffer overnight at 4°C. After washes, embryos were incubated with secondary antibodies diluted in the wash buffer for 2 hours at room temperature with gentle agitation. Staining with Rhodamine Phalloidin (Invitrogen, R415) diluted at 1:500 was performed together with secondary antibodies. Subsequently, embryos were washed in PBST with DAPI (Invitrogen, D3571) at 5 μg/mL and mounted in PBST.

Primary antibodies against GATA4 biotinylated (R&D systems, AF2606), SOX2 (Cell Signaling, 23064), TFAP2C (Cell Signaling, #2320), CDX2 (Biogenex Laboratories, MU392AUC), PARD6B (Santa Cruz Biotechnology, sc-67393), pan-Laminin (Novus Biologicals, NB300-144SS), Collagen IV (Millipore, AB756P), Fibronectin (Proteintech, 15613-1-AP), ITGB1 (Millipore, MAB1997), GFP (chromotek, gb2AF488) were diluted at 1:200. Primary antibodies against active ITGB1 (12G10) (Santa Cruz Biotechnology, sc-59827), ZO1 (Invitrogen, 33-9100), di-phosphorylated myosin regulatory light chain (ppMRLC) (Cell Signaling, 3674), phosphorylated ERM (pERM) (Cell Signaling, 3726) were diluted at 1:100. Primary antibodies against OCT3/4 (Santa Cruz Biotechnology, sc-5279) and KRT8 (Troma-1-C, AB531826) were diluted at 1:50.

The following secondary antibodies were used at 1:400: donkey anti-goat IgG Alexa Fluor 488 (ThermoFisher, A11055), donkey anti-rat IgG Alexa Fluor 488 (ThermoFisher, A21208), donkey antimouse IgG Alexa Fluor 488 (ThermoFisher, A21202), donkey anti-rabbit IgG Alexa Fluor Plus 546 (Invitrogen, A10040), donkey anti-goat IgG Alexa Fluor Plus 555 (Invitrogen, A21432), donkey antimouse IgG Cy5 AffiniPure (Jackson ImmunoResearch, 715-175-150), donkey anti-rabbit IgG 647 (ThermoFisher, A31573).

### Cryosectioning

Pregnant mouse uteri were dissected and handled in KSOM with HEPES. To reduce non-physiological uterine contraction due to the release from connecting tissues, uteri were transferred to pre-warmed 0.5 M MgCl_2_ solution. Uteri were cut into pieces corresponding to the embryo implantation sites, as visually judged by their swollen and opaque appearance under the stereomicroscope. Tissue pieces were immediately fixed in 4% PFA in PBS overnight at 4°C, followed by an overnight wash in PBS at 4°C, and subsequent overnight washes in 12% Sucrose, 15% Sucrose, and 18% Sucrose at 4°C until further use within two weeks. The tissue pieces were dried with KIMTECH paper (Kimberly-Clark) and mixed with M-1 Embedding Matrix for cryosectioning (ThermoScientific, 1310TS). Tissue pieces were mounted and orientated in M-1 Embedding Matrix in Tissue-Tek cryomold (Sakura) and frozen at −80°C.

Cryosectioning was performed with Leica CM3050S cryotome at −16°C, to produce sections of 15-20 μm thickness using low profile microtome blades (Accu-Edge, Sakura). Tissue sections were dried at room temperature, washed in PBST, and permeabilized for 15 min using 0.5% Triton X-100 in PBS. Immunostaining was performed as described above.

### Confocal Imaging

Confocal imaging was performed on Zeiss LSM 780 Confocal Inverted Microscope with LD C-Apochromat 40×/1.1 W Corr objective, using Zen 2012 LSM Black software and LSM880 Airyscan Confocal Inverted Microscope with a C-Apochromat 40x/1.2 NA water immersion objective, using Zen 2.3 SP1 Black software v14.0.0.0. Nuclear immunostaining of OCT3/4, GATA4, and TFAP2C were imaged by LSM780 or LSM880 confocal mode (evaluation of 3E-uterus) with 1 μm Z spacing.

Immunostainings of embryos and tissue sections were also imaged with Airyscan Optimal or Superresolution modes with optimal Z spacing, calculated based on the used imaging settings. The following lasers were used: diode 405 nm, argon multi-line 458/488/514 nm, HeNe 561 nm and 633 nm. Raw Airyscan images were processed by ZEN 2.3 SP1 Black software v14.0.0.0 or v14.0.12.201. See Table S2 for the summary of the microscopy types used throughout the study.

### Light-Sheet Live-Imaging with Muvi-SPIM

#### Microscope and Imaging Settings

The multiview light-sheet microscope is equipped with 2 Olympus 2 mm WD 20x/1.0 NA water immersion objectives (XLUMPLFLN20XW) used for detection, and 2 Nikon 3.5 mm WD 10x/0.3 NA water dipping objectives (CFI Plan Fluor 10X W) used for illumination. The detection path further consists of a filter wheel, a Nikon TI-E 1× tube lens (Nikon Instruments Inc.), and a CMOS camera (ORCA-Flash4.0 V2, Hamamatsu Photonics K.K.), The captured 3D data has a voxel size of 0.295×0.295×1.000 μm^3^ along the X, Y, and Z axes, respectively. The recorded volume size amounts to 302.08×604.15×150-250 μm^3^. The following lasers and filters were used: 488 nm (LuxX^®^ series, Omicron-Laserage Laserprodukte GmbH) and BP525/50 (525/50 BrightLine HC, Semrock, IDEX Health & Science LLC), 561 nm (OBIS LS 561, Coherent Inc.) and LP561 (561 LP Edge Basic Langpass-Filter, Semrock, IDEX Health & Science LLC). Dual light-sheet illumination was used, paired with line-scan detection mode (Medeiros et al., 2015) with a slit width of 40 px. The exposure was set to 30 ms. Live imaging was performed under 5% CO^2^ and 19.5% O^2^ atmospheric conditions at 37 °C inside the controlled environmental imaging chamber.

#### MuVi-SPIM Image Processing

The volumes acquired with the left and right cameras were fused using the Luxendo Image Processor (v2.4.1., Luxendo, Bruker Corp). For further quantification and analysis, the image drift was corrected in Fiji (Schindelin et al., 2012) with the BigDataProcessor2 plug-in (Tischer et al., 2021).

### Light-sheet Live-imaging with InVi-SPIM

An array of micro-cavities was fabricated inside the PEG hydrogel-filled TruLive3D Dishes using custom PDMS stamp, containing a single row of micro-cavities (see “Hydrogel holders and microtopography stamps fabrication”). The dish bottom was covered with 35 μL of the PEG hydrogel precursor mix; the PDMS stamp was carefully placed parallel to the side of the detection objective. After hydrogel solidification for 30-40 min in the incubator, 200-300 μL of PBS was added atop and the stamp was pulled out with forceps. Several washes with IVC1 medium were performed before embryo culture. Embryos were carefully mounted into crypts in a downward mTE orientation, IVC1 medium was added up to 115 μL, and covered with 250 μL mineral oil to prevent evaporation during live imaging. IVC medium was exchanged as described (see “Embryo Culture”). Live imaging was performed under 5% CO^2^ and 19.5% O^2^ atmospheric conditions at 37 °C inside the controlled environmental imaging chamber. The InVi-SPIM is equipped with a Nikon 25x/1.1NA water dipping objective (CFI75 Apochromat 25XC W, Nikon Instruments Inc.) used for detection, Nikon 3.5 mm WD 10x/0.3 NA water dipping objectives (CFI Plan Fluor 10X W) used for illumination, and CMOS camera (ORCA-Flash4.0 V2, Hamamatsu Photonics K.K.). Voxel size: 0.104×0.104×1.000 μm^3^ along the X, Y and Z axes, respectively. The following lasers and filters were used: 488 nm and BP525/50 (525/50 BrightLine HC, Semrock, IDEX Health & Science LLC), 561 nm and LP561 (561 LP Edge Basic Langpass-Filter, Semrock, IDEX Health & Science LLC) Exposure time was set to 50 ms. Imaging was performed with line-scan mode in LuxControl (Luxendo, Bruker Corp).

### Pharmacological Rac1 Inhibition and Live Imaging

Embryos were recovered at E3.5 (D0) and manipulated according to the 3E-uterus protocol. Embryos from the same litter were split into two isolated TruLive3D dish compartments for the parallel live imaging of the treatment and the control conditions. Live imaging started at 30 h counted from the time of embryo recovery. A single mTomato channel was illuminated with a 561 nm laser every 20 min during subsequent live imaging intervals. At 36 h, IVC1 medium in one compartment was exchanged to IVC1 medium supplemented with 100 μM NSC23766 (treatment) and in another compartment to IVC1 medium supplemented with an equal amount of H2O (control). The supplemented medium (for both treatment and control) was exchanged 3-4 times with several-minute incubation time intervals to equilibrate the concentrations. Imaging restarted at 37 h until 48 h. Between 48 h and 49 h, the medium was exchanged in the same way to non-supplemented IVC2 for both the treatment and the control conditions. Live imaging restarted at 49 h and continued until 72 h, after which embryos were fixed and immunostained. Image voxel size: 0.208×0.208×1.000 μm^3^ along the X, Y, and Z axes, respectively.

### Image Analysis Software

Dimension measurements and cell counting were performed with Imaris v9.2.1 (Bitplane). ICY (de Chaumont et al., 2012) was used for cell tracking. Fiji (Berg et al., 2019; Schindelin et al., 2012) was used for kymograph analysis, basal membrane segmentation, contact angle quantification, volume measurements, and fluorescence intensity quantification for plasma membrane proteins. Ilastik (Berg et al., 2019) was used for CDX2-GFP nuclear signal segmentation. Paintera software was used to generate and correct the ground truth segmentation (https://github.com/saalfeldlab/paintera).

### Nuclei Segmentation

The 3D data volumes were acquired with either LSM780 or LSM880 in a confocal mode with a voxel size of 0.207×0.207×1 μm^3^ or 0.23.23×0.23×1 μm^3^, for X, Y, and Z dimensions, respectively. The channels corresponding to anti-OCT3/4, anti-GATA4, and anti-CDX2 immunostainings were used.

A 3D UNet (Çiçek et al., 2016) was trained with a multi-task objective: predicting the binary nuclei mask in the first output channel and predicting the nuclei boundaries/outlines in the second output channel. The boundary predictions were then used to recover the individual nuclei using PlantSeg’s ‘MutexWS’ partitioning algorithm. The nuclei foreground prediction is used in post-processing for removing spurious instances in the background.

Model training was performed iteratively with an increasing amount of ground truth data. Starting from four initial ground truth data volumes, in each iteration, we trained the network, performed the segmentation, and manually proofread the results in order to increase the training set and accuracy. In total, 22 training and 13 validation data volumes were used for the final model training. The size of the training volumes ranged from [117, 703, 377] to [162, 1052, 1840] voxels in X, Y, and Z dimensions.

### Membrane-based Cell Segmentation

The data volumes were acquired with MuVi-SPIM (see “Light-Sheet Live-Imaging with Muvi-SPIM”). A dedicated 3D UNet was trained to predict the foreground membrane mask, which was used for the final cell segmentation with PlantSeg’s ‘GASP’ agglomeration algorithm. The ground truth for the network training was bootstrapped by initially segmenting the stacks with pre-trained PlantSeg models (‘confocal_unet_bce_dice_ds2x’), followed by manual correction of the erroneous cells. In total, four annotated stacks were used for training and one for validating the network. Both nuclei and membrane UNets were trained using Adam optimizer (Kingma and Ba, 2014) with β1=0.9, β2=0.999, L2 penalty of 0.00001, and initial learning rate ε=0.0002. Networks were trained until convergence for 100K iterations, using the PyTorch framework (Paszke et al., 2019). The models with the best score on the validation set were selected.

### Embryo Staging by Cell Numbers

Cell counts for E4.5-E6.0 *in utero* embryos were obtained from the previous study (Ichikawa et al., 2022). Cells for E3.5 *in utero* embryos were manually counted based on GATA4 and SOX2 immunostaining. Linear regression analysis for embryo staging was performed as described (Ichikawa et al., 2022). For successfully developed 3E-uterus embryos, epiblast (EPI) cells were defined based on the nuclear OCT3/4 expression. Cells with nuclear GATA4 expression overlying epiblast cells were defined as visceral endoderm (VE). OCT3/4 and GATA4 channels were used for automatic EPI and VE nuclei segmentation (see “Machine-learning-based nuclei segmentation”). For the absolute quantification accuracy, manual correction and cell counting were performed on top of the automated nuclei segmentation.

### Evaluation of 3E-uterus Efficiency

Efficiency was quantified as a percentage of successfully developed embryos among all embryos at day 3 of 3E-uterus. 3E-uterus embryo were classified as successfully developed if three criteria were met (Figure S1E):

I. Egg cylinder formation, defined as EPI tissue located within a VE layer with the basal membrane in between.
II. Alignment of the egg cylinder axis with the crypt axis. The embryos with an evident upward egg cylinder orientation were excluded from quantifications due to an experimental error of embryo positioning (corresponding to less than 5% of samples).
III. Formation of the Reichert’s membrane, determined as a basal membrane underneath TB which, at the top of the egg cylinder, was required to continue into the basal membrane between EPI and VE. To directly assess the criteria I-III, the simultaneous immunostaining against OCT3/4, GATA4, Collagen IV. or pan-Laminin, and nuclei (DAPI), was performed each time. For evaluation of 3E-uterus efficiency, three independent experiments were performed, among which 46% of embryos (12 of 26) met all the above-mentioned criteria.

#### Efficiencies for crypt diameter evaluation (Figure S1F) were calculated as follows

80 μm crypt diameter: 0/4, 0/5, and 1/8 (the number of successfully developed embryos divided by the total number of embryos); three independent experiments.

100 μm: 0/3, 2/9, and 2/8; three independent experiments.

120 μm: 1/3, 1/5, and 2/5; three independent experiments.

140 μm: 3/8, 2/5, 1/3, 2/7, 1/4; five independent experiments.

160 μm: 2/4, 0/5, 1/5; two independent experiments.

#### Efficiencies for the hydrogel stiffness evaluation (Figure S1H) were calculated as follows

1.5% PEG-PEP content: 3/8, 3/8, and 2/8 (the number of successfully developed embryos divided by the total number of embryos); three independent experiments.

1.75%: 3/9, 5/8, 2/7, and 4/8; four independent experiments.

2%: 2/7, 3/6, and 4/6; three independent experiments.

2.25%: 0/6, 6/13, and 1/5; three independent experiments.

2.5%: 1/6, 2/7, and 4/17; three independent experiments.

2.75%: 1/9, 5/15, and 1/9; three independent experiments.

6%: 1/6, 0/7; two independent experiments.

7%: 1/11, 0/5; two independent experiments.

### Extraembryonic Ectoderm Cell Number Counting

Nuclei were counted based on CDX2-GFP signal in MuVi-SPIM 3D data volumes using automated Spots detection with manual correction in Imaris.

### Trophoblast Cell Tracking

Individual cells on the mural TE side of the H2B-GFP expressing embryos were tracked in 3D over 18-24 hours of imaging, starting from 30 hours post E3.5 recovery.

### Cell Speed Quantification

The mTE/TB cell speed was quantified as the Euclidian distance between the mTE/TB nuclei positions in the adjacent hours of live imaging using a sliding time window with a size corresponding to one hour and a step size of an image time resolution (10 or 15 min) (Figure S2D).

### Cell Directionality Quantification

The sliding window (see “Cell Speed Quantification”) was used to define mTE/TB vector between time points. We calculated the angle (a) between mTE/TB vector and the unit vectors corresponding to X, Y, and Z axes. ‘Directionality’ was calculated as (180 – α)/90 – 1, ranging from −1 to 1 values (Figure 2I, Figure S2C).

### Quantification of the neighbourhood persistence

The cell neighborhood was defined for each TB cell as the nearest four TB cells. Persistence of neighborhood was quantified as a proportion of cell neighbors maintained between adjacent hours of live imaging, ranging from 0 to 1.

### Fluorescence Intensity Quantification for Plasma Membrane Proteins

Identical imaging settings were applied for the samples in Figure 3A-D to enable comparison. The fluorescence signal of ZO1, PARD6B, and phospho-Ezrin/Radixin/Moesin was measured in Fiji using a line tool, 5 pixels in width, drawn along the cell’s perimeter. Signal intensity values along the cell perimeter were exported for analysis and visualization in R. The signal was normalized to the average nuclear DAPI signal within the same Z plane.

### Quantification of the Contact Angle at the Mural TE-hydrogel Interface

Image volumes were manually transformed with BigDataProcessor2 Fiji plugin for vertical crypt alignment along the y-axis. Images were XZ-resliced followed by 180° radial reslice about the center of the line of symmetry. Microwell surface was identified based on the background hydrogel fluorescence. Fiji’s Ange tool with a handle length of 15-20 μm was used to quantify the angle (θ) between the crypt surface and the cell membrane on the mural and polar TE sides. θ values were quantified on the left and right sides of the image every 30°. The final θ value represents the averaged value across the crypt circumference.

### Kymograph Analysis

To quantify mural TE and EPI displacements, a kymograph was drawn parallel to the crypt axis and the edge of the membrane signal was tracked. Per each embryo, the values were averaged across three lines per Z-slice in three different Z locations.

### Polar TE Cell Shape Analysis

Polar TE cell length and the width were manually measured with Imaris based on the overlay of the cell membrane segmentation output and the raw signal. The dimensions were measured for 15-20 polar TE cells per embryo every hour of live imaging.

### Embryo Length Analysis

Embryo length was quantified in 3D as a distance between the outermost giant trophoblast nucleus and the outermost nucleus of the polar TE/ExE along the crypt axis (Figure 3H).

### Basal Membrane Segmentation

Segmentation of the basal membrane (BM) between EPI and PrE from the Reichert’s membrane was performed with the segmentation editor (https://imagej.net/plugins/segmentation-editor) in Fiji based on anti-Collagen IV or anti-pan-Laminin immunostaining data. 2D Roi with the BM data signal were converted into continuous contours using a custom Python script.

### Middle Axis Estimation and Length Computation

The binary 3D segmentation of the basal membrane (BM) between EPI and PrE was used for analysis. First, the Euclidean Distance Transform (DT) was applied to the 3D segmentation to construct a directed graph in which the nodes are the non-zero valued pixels of the DT. The edges of the graph were assigned with weights that represent the difference between the global maximum of the DT values and the DT value of the target node. The shortest path in the weighted graph was then computed between two nodes that correspond to manually annotated points on the specimen’s surface that mark its extreme poles (Dijkstra, 1959). The nodes on the shortest path were used to fit an open cubic B-spline (Schoenberg, 1969) curve that approximates the middle axis. Finally, the integration over the spline was performed in order to obtain the arc-length of the egg cylinder.

### Diameter Estimation

Similarly, the binary 3D segmentation of the BM was used for diameter estimation. Two landmark points, which correspond to the middle of the EPI tissue density, were manually annotated on the specimen’s surface. The landmark points were then used to determine a plane that intersects the specimen orthogonally with respect to the estimated middle axis (see the previous section). More specifically, the plane was fitted such that it minimizes the Euclidean distance to the landmark points under the constraint of being orthogonal to the middle axis. The closed circular curve, resulting from the intersection of the specimen and the plane, was then used to compute the diameter.

## ACKNOWLEDGMENTS

We would like to thank all members of the Hiiragi group for discussions, comments, and critical reading of the manuscript; Lidia Perez, Ramona Bloehs, and Stefanie Friese for their technical support; former interns Nisha Veits and Falk Farkas for their technical support; Qin Yu from A. Kreshuk group for the help with image segmentation; all members of the Erzberger group for their input on theoretical analysis. We are thankful to the EMBL animal facility for the mouse work; Christian Kieser and Tim Hettinger at the EMBL electronic and mechanical workshops for their technical support; EMBL Advanced Light Microscopy Facility and Christian Tisher for the help with image analysis; Luxendo for their technical support with light-sheet microscopy. We would like to thank Dr. Terry Lechler for the ZO1-GFP mouse line and Dr. Isabelle Migeotte for the Rac1^flox^ mouse line. V.B. is supported by the Add-on Fellowships for Interdisciplinary Life Science of the Joachim Herz Stiftung (JHS). The Hiiragi laboratory is supported by EMBL, Hubrecht Institute, and the European Research Council (ERC Advanced Grant “SelforganisingEmbryo”, grant agreement 742732).

## AUTHOR CONTRIBUTIONS

V.B., M.N., M.L., and T.H. conceived and designed the project; V.B. developed protocols and conducted experiments and analysis. M.N. manufactured hydrogels, stamps, and helped with the hydrogel preparation. V.B. and D.K. developed sample mounting for MuVi-SPIM; D.K. developed incubation and helped with MuVi-SPIM live imaging and processing. A.W. and V.B. developed and trained the image segmentation models. R.B. and A.E. conceived and conducted the theoretical analysis. S.R. synthesized the hydrogels. J. H. performed embryo shape analysis. V.B. prepared the figures and movies. V.B., A.E., and T.H. wrote the manuscript with inputs from all authors.

## Supplemental Note

### 1 Droplet wetting model of embryo implantation

A thermodynamic theory of capillary phenomena can be formulated by using the free energy *H* of an incompressible liquid droplet [3, Sec. 5.6]. Considering the whole embryo as such a droplet and the 3E-uterus as a solid substrate, we then pose

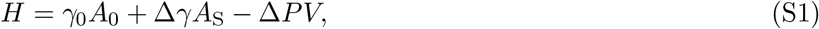

in which *γ*_0_ is the surface tension between the medium and the embryo, Δ*γ* = *γ*_E_ — *γ*_M_ is the Young tension—the difference between the surface tension of embryo-substrate (*γ*_E_) and medium-substrate (*γ*_M_) contacts,— and *A*_0_ and *A*_S_ are the areas of the embryo-medium and embryo-substrate contacts respectively (Fig. 4D). The Laplace pressure Δ*P* acts as a Lagrange multiplier to the volume of the embryo *V* = const.

#### 1.1 Equilibrium solutions in cylindrical geometry

First we consider the simplest limiting case of an embryo within a cylindrical confinement and seek the equilibrium solutions for the droplet shape. Given uniform interfacial tensions, the droplet must manifest cylindrical symmetry (Fig. S4A). For positive *γ*_0_, the droplet-medium interface therefore takes the minimal surface corresponding to a spherical cap. We denote by *h* the distance between the contact lines, by *y* the height of the spherical caps, and by *r* the radius of the cylinder. When the height *y* is negative, the spherical caps curve inwards into the embryo. With these definitions, the contact areas and volume are given by

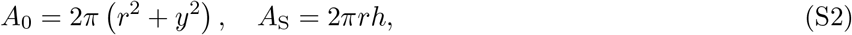

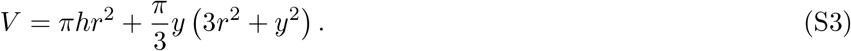

The value of *h* is determined by the volume constraint

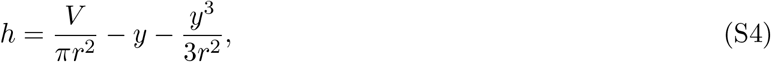

and the equilibrium conditions for *h* and *y* read

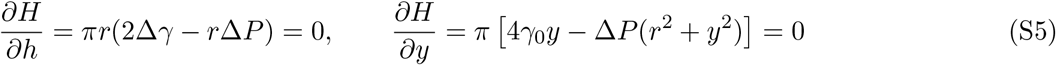

vhich recapitulate the Laplace law for the pressure Δ*P* = 2Δ*γ/r* [3, Sec. 5.6] and yield

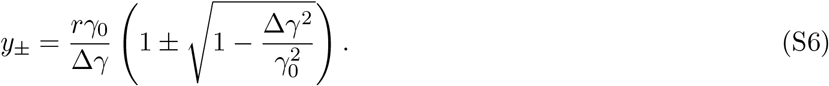

Equilibrium solutions exist only for the partial wetting regime —1 < Δ*γ*/*γ*_0_ < 1 (Fig. S4C). For Δ*γ*/*γ*_0_ < –1, the droplet spreads along the surface of the confinement completely (total wetting), whereas for Δ*γ*/*γ*_0_ > 1, an interface between the droplet and the substrate is not energetically favored (dewetting) and therefore a droplet with a volume below the confinement limit detaches from the substrate and takes on a spherical shape (Fig. S4D). In the partial wetting regime, Eq. (S6) has a stable and an unstable solution *y*_-_ and *y*_+_ respectively, for which

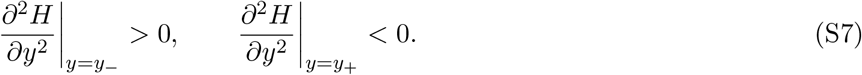

The contact angle *θ* = *φ* + *π*/2 is related to the angle of the spherical cap *φ* and therefore

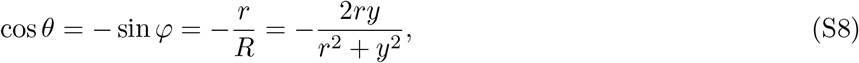

in which we used the formula for the radius of curvature of the spherical cap *R* = (*r*^2^ + *y*^2^)/(2*y*). If we consider only the stable solution in Eq. (S6) we further obtain

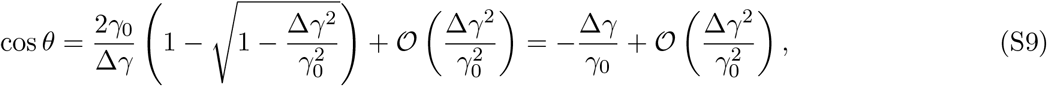

where the last equality recapitulates the Young law for the wetting angle [1, Sec. II.A.1].

#### 1.2 Shape dynamics in frustum geometry

To model the changes in trophoblast adhesion during implantation, we now consider a dynamic decrease of the interfacial tension between the embryo and the 3E-uterus, and calculate the resulting shape dynamics by taking into account the broken up-down symmetry of the conical-frustum confinement. We assume that the contact angle remains close to its equilibrium value as the adhesion of the embryo to the 3E-uterus changes.

We describe the system’s state by four thermodynamic variables ***z*** = (*z*_1_, *z*_2_, *z*_3_, *z*_4_) (Fig. S4B): positions of the top and bottom contact lines *z*_1_ and *z*_2_, respectively, and the heights of the top and bottom spherical caps *z*_3_ and *z*_4_. The radii of the frustum’s horizontal sections through the contact lines are then *r_a_* = *χz*_1_ and *r*_b_ = *χz*_2_, in which *χ* = tan(*α*/2) with α the conical angle.

With these definitions we obtain the following expressions for the contact areas and the volume of the droplet:

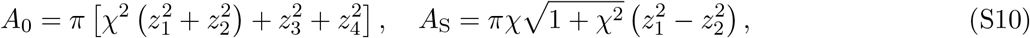

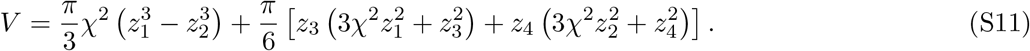

The above description yields the cylindrical geometry *r_a_* → *r, r_b_* → *r* as we take the limit *χ* → 0.

From the equilibrium condition for the frustum geometry (*∂H/∂z*_*i*=1,2,3,4_ = 0) we can derive the following formulas for the Laplace pressure and the Young tension:

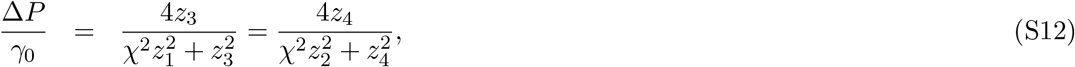

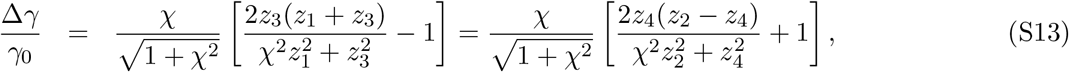

which should hold independently for the upper and lower spherical caps described by equilibrium values of (*z*_1_, *z*_3_) and (*z*_2_,*z*_4_) respectively (Fig. S4B).

Equations (S12) and (S13) applied to the experimentally measured geometry of the embryos (Sec. 2) yielded different results for the upper and lower caps, (*z*_1_,*z*_3_) and (*z*_2_,*z*_4_) respectively, which are thus inconsistent with the equilibrium state. Therefore we concluded that the observed series of *z*(*t*) outline a sequence of nonequilibrium states as discussed shortly below.

Assuming a linear constitutive relation between the energy gradient and the velocities of the variables *z*_*i*=i,2,3,4_ with dissipative coefficients *λ*_*i*=1,2,3,4_, we obtain equations of motion

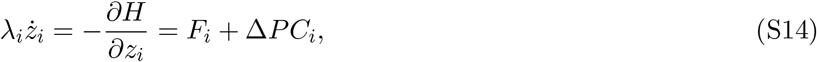

in which

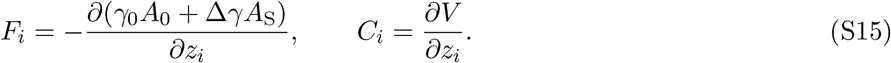

The Young tension Δ*γ*(*t*) is now a time-dependent active parameter, describing the increase in adhesion between the embryo and the substrate. We describe the dependence of the Young tension on time t by a generic sigmoid shape (Fig. S4E):

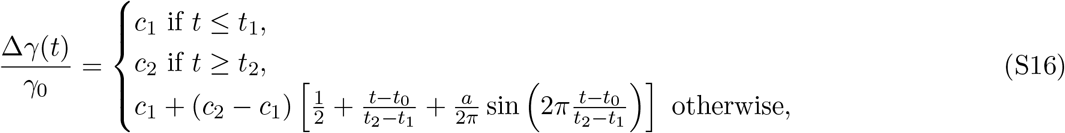

in which *c*_1_ and *c*_2_ are constant values, *t*_1_ and *t*_2_ are respectively the beginning and the end of the adhesion decrease with the mid time *t*_0_ = (*t*_1_ + *t*_2_), whereas *α* ∈ [-1,1] is a modulation amplitude.

Furthermore, during the implantation process the embryo also regulates its volume, which can be incorporated into our model by making the volume constraint time-dependent:

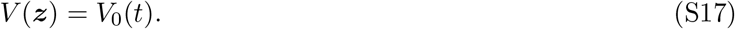

The value of the Lagrange multiplier Δ*P* can be determined by differentiating the constraint Eq. (S17),

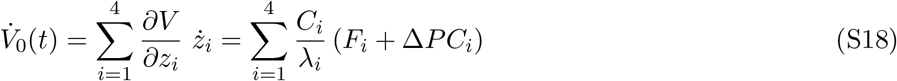

which is solved by

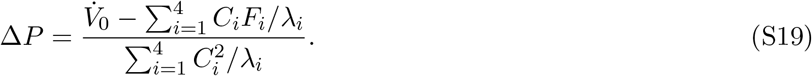

Given an initial condition ***z***(0) and the functions Δ*γ*(*t*) and 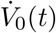, the equations of motion (S14) can be integrated for ***z***(*t*).

In the course of motion defined by Eq. (S14) the free energy changes as

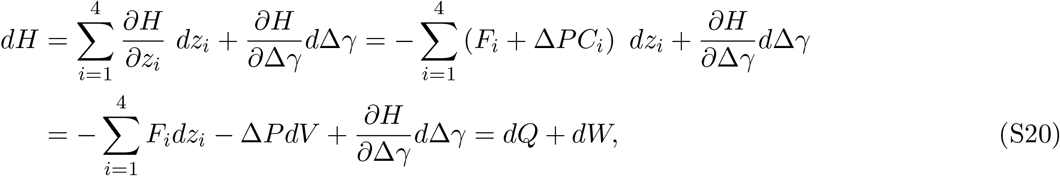

which recapitulates the first law of thermodynamics if we identify the heat and the active work, respectively,

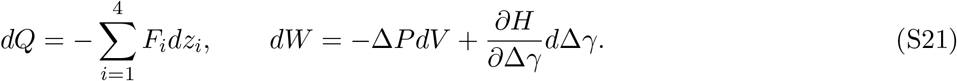

The two active contributions of the work *dW* = *dW*_1_ + *dW*_2_ correspond to the volume change and the adhesion change:

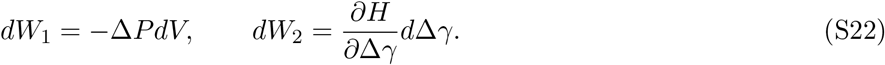

Note that, when the volume *V* is conserved, i.e. *V*_0_(*t*) ≡ const, the forces of constraint remain perpendicular to the system’s trajectory ***dz***(*t*) and thus do no work (*dW*_1_ = 0). This observation follows from the geometric analysis of Eqs. (S14)–(S19), which are entirely analogous to the isokinetic thermostat in molecular dynamics as discussed by Evans and Morriss [2, Sec. 5.2].

Given that the dissipative coefficients λ_*i*_ are large compared to the speed of the adhesion change |*c*_2_ – *c*_1_|/(*t*_2_ – *t*_1_) (Fig. S4E), the system relaxes slowly in the response to the Young tension change and, thus, the observed time series of ***z***(*t*) may correspond to transient states substantially far from equilibrium.

### 2 Comparison with experimental data

Time series were acquired with a time resolution of 1 hour for the whole-embryo volume *V*_0_, as well as for the contact angles, *θ_a_*(*ω_i_*) and *θ_b_*(*ω_i_*), and positions of the top and bottom contact lines with respect to *z*_0_, *x_α_*(*ω_i_*) and *x_b_*(*ω_i_*), at several points *ω*_*i*=1, 2..._ around the conical axis. The heights of the spherical caps were estimated as

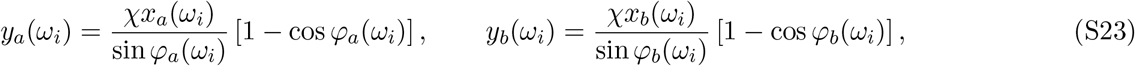

in which

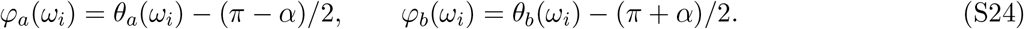

By averaging the above measurements over the points *ω_i_* we find the time series for the thermodynamic variables of interest

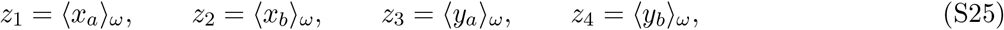

and their standard deviations *σ*_*i*=1,2,3,4_.

A smooth representation of the embryo volume was constructed by interpolating the experimentally mea-sured volume with

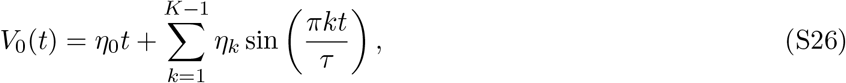

in which *τ* is the total observation time, and *K* is the number of timepoints, which yields a derivative with a spectral accuracy [5, Chapter 4]

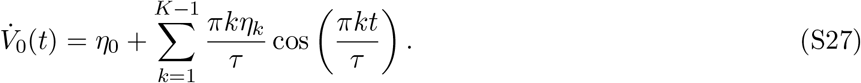

The frustum-angle tangent is *χ* = 0.1, as measured from the imaging data.

#### 2.1 Simulation-based inference

To determine the dissipative coefficients λ_*i*=1,2,3,4_ and parameters of the Young tension Δ*γ*(*t*) given by Eq. (S16) we used simulation-based inference [4] with the time series of ***z***(*t*), the standard deviations ***σ***(*t*), and the volume derivative 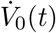, which were acquired from the experiments as described above. Furthermore we assume λ_1_ = λ_2_, λ_3_ = λ_4_, and |Δ*γ*| ≤ *γ*_0_. Because time-series of the geometric data do not provide complete information about quantities involving energy and mass, we adopt a custom system of physical units based on hours for time, *μ*m for length, and *γ*_0_ for tension.

In total we have seven fitting parameters: λ_1_, λ_3_, *c*_1_, *c*_2_, *t*_0_, Δ*t* = *t*_2_ — *t*_1_, and *a* (Table S3). Assuming uniform prior distributions of these parameters, we applied two rounds of sequential neural posterior estimation [4] with 10^6^ simulations in each round and Gaussian kernel-mixture representation of the probability density. The neural posterior estimator was trained directly on the time series ***z***(*t*) generated by Eq. (S14) with the experimentally determined 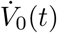 and a superimposed random Gaussian error of the zero mean and the observed standard deviation ***σ***(*t*). The parameter values thus estimated are mostly consistent across the three embryos for which we have measurements, with a somewhat larger variability of the dissipative coefficients λ_*i*_.

## Notes

https://github.com/kreshuklab/mouse-embryo-seg

https://doi.org/10.5281/zenodo.6546550

### Competing Interest Statement

The authors have declared no competing interest.

